# Reduced Microtubule Polyglutamylation Perturbs Neuronal Function and Accelerates Tau-Induced Neurodegeneration

**DOI:** 10.64898/2026.07.31.742027

**Authors:** Shreyangi Chakraborty, Silvia Poggini, Michaela Rusková, Isabelle Lavenir, Davide Boido, Adela Karhanova, Laetitia Besse, Naomi Ciano Albanese, Yana Al-Inaya, Laura Lebrun, Veronique Henriot, Zdenek Lansky, Michel Goedert, Luisa Ciobanu, Martin Balastik, Igor Branchi, Carsten Janke, Maria M. Magiera

## Abstract

Microtubules are essential for neuronal functions and are involved in many neurodegenerative disorders. A posttranslational modification (PTM) of neuronal microtubules, polyglutamylation, was shown to control cargo transport, Tau protein binding to microtubules and to induce early-onset neurodegeneration when exceeding physiological levels. To determine the physiological importance of this PTM, we characterised mouse models with reduced polyglutamylation. We demonstrate that these mice develop late-onset atrophy of the olfactory bulb and frontal cortex, progressively reduced dendritic spine density, and deficits in olfaction, memory, and cognitive control. This indicates an important role of polyglutamylation in neuronal function. To test whether the observed defects are linked to neurodegeneration, we combined mice with reduced polyglutamylation with a transgenic model for tauopathy. Strikingly, reduced polyglutamylation accelerated the premature demise of mice transgenic for human Tau carrying the P301S mutation. Our work thus reveals key roles of microtubule polyglutamylation in neuronal function with implications for late-onset neurodegeneration linked to Tau pathology.

## Introduction

Microtubules, as key components of the cytoskeleton, are essential for virtually every eukaryotic cell type. In neurons, which are morphologically and functionally highly complex, microtubules play essential roles in neuronal development and functional maintenance throughout life^1–4^. A variety neurodegenerative disorders are associated with dysfunctions of the microtubule cytoskeleton; however, in many cases, the molecular mechanisms causing the cascade of pathological processes are still not fully understood^5–8^.

A striking feature of microtubules is their extraordinary degree of conservation throughout evolution – microtubules, and their building blocks, the tubulins, are virtually identical from protists to complex differentiated cells in humans^9^. It is thus a long-standing and baffling question how microtubules can adapt to so many different functions, and how these functions can be coordinated in a single cell. As part of the answer emerges the tubulin code, a combination of molecular mechanisms to diversify tubulins by expression of alternative gene products, called tubulin isotypes, or by a plethora of posttranslational modifications (PTMs) of tubulin^10, 11^.

Tubulin PTMs were discovered decades ago, but for a long time could not be functionally characterised, which is why they remain relatively underexplored. Recent years have seen a surge of research on tubulin PTMs fuelled by the discovery of the enzymes^12–24^ and the generation of animal models^16, 25–31^ now allowing to reveal their physiological functions. Many of the currently known PTMs are enriched in the nervous system, i.e., in neurons. Polyglutamylation, a PTM generating branched peptide chains made of glutamates^32–34^, becomes strongly enriched during neuronal development^35, 36^, and most of the enzymes catalysing it show continuous expression into adulthood^37^. In contrast to another PTM, detyrosination, which has deleterious effects on neuronal development when upregulated^26^, polyglutamylation has less essential, though important, developmental roles^38^. However, upregulation of polyglutamylation (hyperglutamylation) in mice with mutations in the deglutamylase CCP1 causes early-onset neurodegeneration with a characteristic degeneration of the cerebellar Purkinje cells^16, 29^, which is not the case when tyrosination is abolished in these cells^39^. Combinatory mouse models revealed that the accumulation of polyglutamylation generated by the neuronal polyglutamylase TTLL1^29^, and partially also by TTLL4^40^, is causative for this degeneration. Purkinje cell degeneration could be entirely prevented by deletion of *Ttll1* and did not occur over the entire lifetime of the animals^29, 40, 41^. The clinical relevance of this discovery was concomitantly revealed by the discovery of a human condition caused by inactivating mutations of CCP1, whose symptoms were phenocopied by the corresponding knockout mice to a surprising extent^42^, and of which a growing number of cases are being identified^43–46^.

At the molecular level, polyglutamylation controls cargo transport^29, 37, 41, 47^, the interactions of a wide variety of microtubule-associated proteins (MAPs)^48, 49^ as well as enzymatic microtubule severing^50–52^. Tight control of these microtubule-based processes is particularly crucial for neurons because these cells are exceptionally long-lived, can have extensions that are extraordinarily ramified (e.g. dendrites of the cerebellar Purkinje cells^53^) or more than one meter long (axons of peripheral neurons). Intriguingly, perturbations of all the molecular microtubule-based events (transport, binding of MAPs and severing) have been linked to some neurodegenerative disorders^4, 8, 54–56^. Given that polyglutamylation affects all of them, and that hyperglutamylation leads to massive neurodegeneration, it is perceivable that precisely controlled polyglutamylation levels on microtubules are essential for neuronal homeostasis. Here we address this question by examining mice with strongly reduced polyglutamylation, focusing on mice lacking the neuronal polyglutamylase Ttll1.

Unlike *Ccp1^-/-^* mice with excessive polyglutamylation, *Ttll1^-/-^* mice show no signs of early-onset neurodegeneration. Observation of brain morphology of ageing mice, however, revealed progressive atrophy of the olfactory bulb and the frontal part of the cortex, accompanied by reduced numbers of neurons, indicative of neuronal loss. We also found a progressive reduction of dendritic spine density, as well as deficits in olfaction, memory, and cognitive control, indicating the presence of early neuronal defects that over time can lead to neuronal dysfunction. Given that polyglutamylation modulates the interaction of microtubules with the protein Tau^49^, which, beyond its role as MAP in healthy neurons, is also responsible for neurodegeneration in tauopathies^57, 58^, we also tested the impact of microtubule polyglutamylation on Tau pathology. Decreasing polyglutamylation in the Tau^P301S^ model for tauopathy^59^ accelerated the demise of these mice, demonstrating a functional link between polyglutamylation and Tau dysfunction. We thus demonstrate a key role of polyglutamylation in neuronal function and homeostasis and provide a mechanistic link between this tubulin PTM and Tau-related neuropathologies.

## Results

### Knockout of TTLL1 and TTLL7 reduces tubulin polyglutamylation in the mouse brain

To address the importance of tubulin polyglutamylation in maintaining neuronal homeostasis, we used mice lacking the two main polyglutamylases expressed in the brain, TTLL1 and TTLL7. To evaluate the tubulin PTM landscape in the brain after knocking out these two genes, brain extracts from wild-type, *Ttll1^-/-^*, *Ttll7^-/-^* and *Ttll1^-/-^Ttll7^-/-^*mice were run on SDS-PAGE gels that allow separation of α- and β-tubulin^60^ and probed with antibodies for different tubulin PTMs (Fig. 1a,c). To relate the level of each PTM to total tubulin levels, the samples were first adjusted to the similar amounts of total α-tubulin with 12G10 and AXO45 antibodies. We then assessed the levels of different tubulin PTMs using both, previously validated and novel antibodies (Fig. 1c).

**Figure 1:**
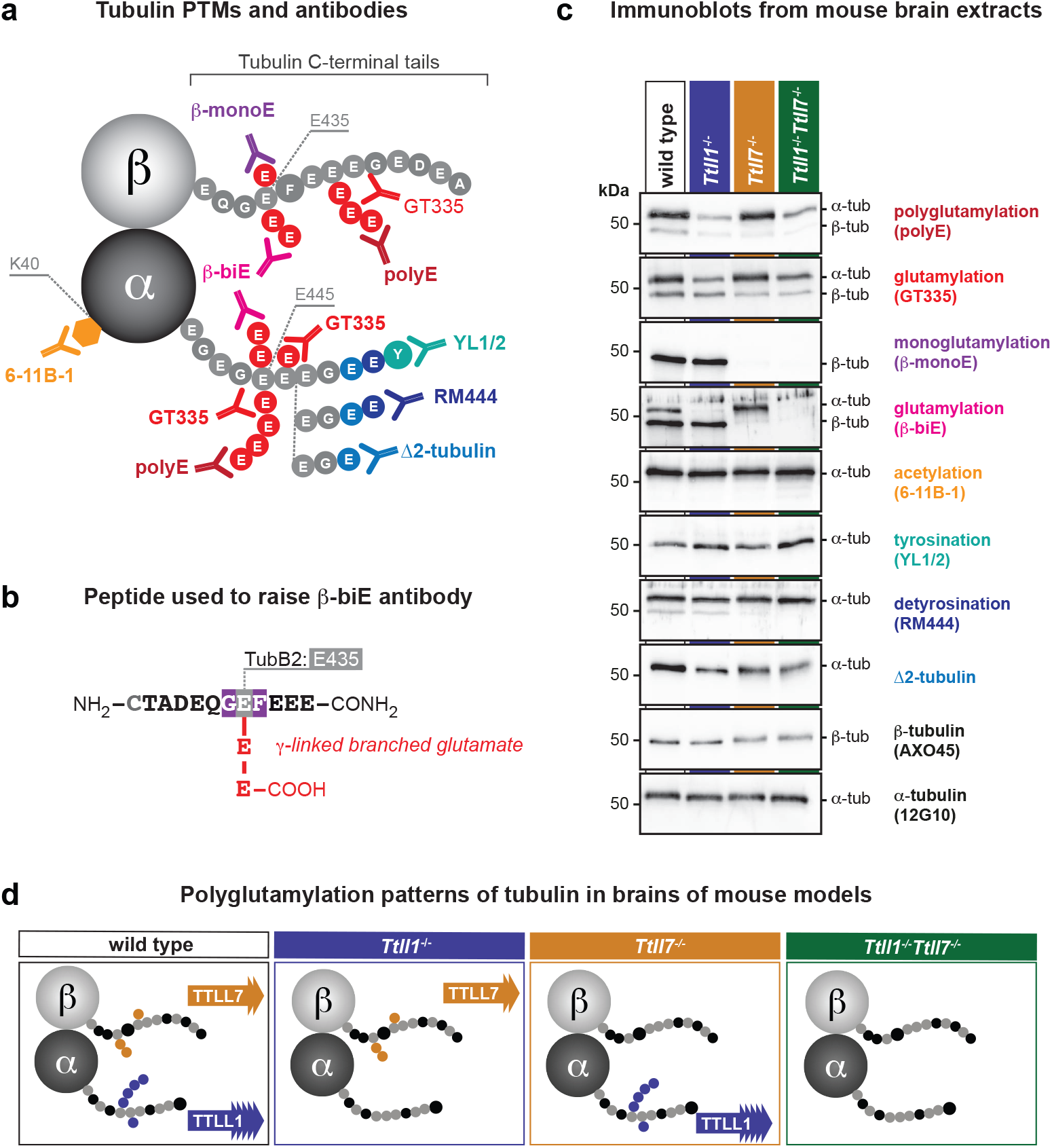
Characterisation of tubulin PTMs in mice lacking TTLL1 and TTLL7 glutamylases. **a)** Schematic representation of posttranslationally modified C-terminal tails of α- and β- tubulins showing epitopes recognised by polyglutamylation-specific antibodies: polyE, GT335, β-monoE, and β-biE, as well as acetylation (6-11B-1), tyrosination (YL1/2), detyrosination (RM444) and Δ2-tubulin. **b)** Synthetic peptide used to generate the β-biE antibody. The β2-tubulin peptide encompassing E435 is shown, with acidic (grey), neutral (black), and aromatic (purple) residues. A γ-linked glutamate branch (red) was introduced to mimic posttranslational bi-glutamylation. **c)** Immunoblot analysis of brain extracts from 2-month-old wild-type, *Ttll1*^-/-^, *Ttll7*^-/-^, and *Ttll1*^-/-^*Ttll7*^-/-^ mice probed with polyE, GT335, β-monoE, and β-biE antibodies to detect different subtypes of (poly)glutamylation, antibodies for acetylation (6-11B-1), tyrosination (YL1/2), detyrosination (RM444), Δ2-tubulin. Total tubulin levels were determined with the pan-β-tubulin (AXO45) and pan-α-tubulin (12G10) antibodies. Molecular weight markers are indicated on the left (kDa), position of α- and β-tubulin bands are indicated on the right. **d)** Schematic representation of polyglutamylation patterns of tubulin found in mouse models used in this study. TTLL1 generates short and long glutamate chains on α-tubulin; TTLL7 generates glutamylation on β-tubulin.

Polyglutamylation was tested using GT335, which recognises the branch point of glutamate chains^61^, polyE detecting C-terminal glutamate chains of at least 3 residues (regardless of the substrate of polyglutamylation)^16, 62^ and β-monoE specifically detecting a glutamate branched on E435 of β2-tubulin (TubB2)^41^ (Fig. 1a). We also included a new polyclonal antibody, β-biE, raised against the bi-glutamylated E435 residue of β2-tubulin (Fig. 1b), but also detecting glutamylated α-tubulin, making it a good tool to assess glutamylation of both α- and β-tubulin (Fig. 1c).

In extracts from wild-type brain, GT335 and polyE strongly labelled α-tubulin and showed a faint signal on β-tubulin; β-monoE labelled β-tubulin only, and β-biE detected strong signals on both α- and β-tubulin (Fig. 1c; S1). In *Ttll1^-/-^* brain extracts, the labelling of α-tubulin by polyE, GT335 and β-biE was almost entirely absent, whereas β-monoE and β-biE antibodies detected glutamylation levels on β-tubulin similar to wild-type (Fig. 1c; S1). In *Ttll7^-/-^*brain extract, polyE, GT335 and β-biE labelling remained unchanged on α-tubulin, while β-monoE and β-biE labelling of β-tubulin was absent (Fig. 1c; S1). In *Ttll1^-/-^Ttll7^-/-^* brain extract all glutamylation antibodies showed faint or no labelling of α- and β-tubulin (Fig. 1c; S1). These observations confirmed previous observations that TTLL1 preferentially modifies α-tubulin while TTLL7 selectively targets β-tubulin^41, 49^. We show now that in the absence of both enzymes, polyglutamylation levels on α- and β-tubulin are reduced to the same extent as in the single knockouts – i.e. there is almost no tubulin polyglutamylation detectable at all – implying that TTLL1 and TTLL7 generate the majority of polyglutamylation in the brain, and, most importantly, their loss cannot be compensated for by other TTLL enzymes (Fig. 1d).

We also investigated whether other tubulin PTMs are affected in mice lacking TTLL1, TTLL7, or both enzymes. We tested tubulin acetylation, tyrosination, detyrosination and Δ2-tubulin. We observed no significant alterations of those PTMs across genotypes (Fig. 1c), highlighting the specificity of glutamylases to tubulin glutamylation, and the lack of detectable crosstalk with other tubulin PTMs in the context of the brain. The loss of polyglutamylation specific to the two modifying enzymes was recently also confirmed on purified tubulin from brain tissue of mice used here by mass spectrometry^63^. In conclusion, TTLL1 and TTLL7 show specific activities generating either α- or β-tubulin glutamylation, and their knockout does not critically affect the levels of other tubulin PTMs (Fig. 1c,d).

### Reduced tubulin polyglutamylation causes late-onset morphological defects in the brain

To determine the impact of changed polyglutamylation patterns in the brains of our mouse models, we first assessed gross morphology of wild-type, *Ttll1^-/-^*, *Ttll7^-/-^*, and *Ttll1^-/-^Ttll7^-/-^*brains at 2 months, 1 year, and 2 years. Macroscopic observation of fixed brains showed a reduced size of the olfactory bulb and a potential atrophy in the frontal part of the cortex of *Ttll1^-/-^* and *Ttll1^-/-^Ttll7^-/-^*, but not *Ttll7^-/-^* brains. These changes were not visible at 2 months, became apparent at 1 year, and persisted to 2 years (Fig. 2a; S2), suggesting that loss of TTLL1-generated polyglutamylation leads to structural defects in the brain that are potentially causing degeneration. While degeneration also occurs in brains with hyperglutamylation caused by the loss of Ccp1 deglutamylase^16, 29^, the timeline of changes and the affected brain regions and cell types was different, indicating that underlying mechanisms might not be identical.

**Figure 2:**
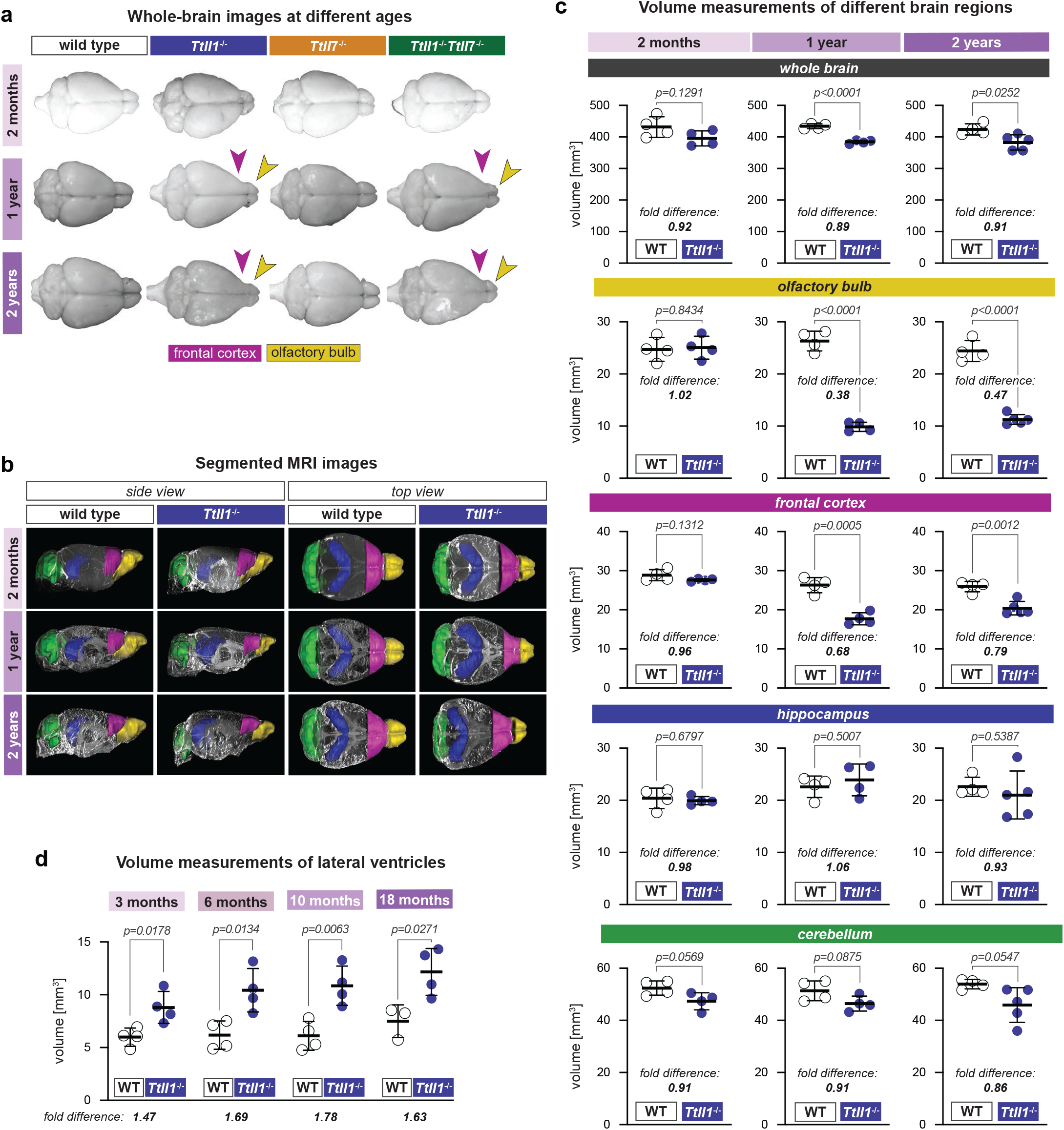
Volumetric analysis of brains of polyglutamylation-deficient mice. **a)** Representative macroscopic images of brains from wild-type, *Ttll1*^-/-^, *Ttll7^-/-^*, and *Ttll1*^-/-^ *Ttll7*^-/-^ mice at 2 months, 1 year, and 2 years. Images from Fig. S2. **b)** 3D-segmented MRI reconstructions of wild-type and *Ttll1^-/-^* brains at 2 months, 1 year, and 2 years shown in side view (left) and top view (right) using Imaris. Colour coding indicates different brain regions analysed: olfactory bulb (yellow), frontal cortex (pink), hippocampus (purple), and cerebellum (green). Images from Fig. S3. **c)** Quantification of volumes of segmented brain regions (from **b**) in wild-type (open circles) and *Ttll1*^-/-^ (blue filled circles) mice at 2 months, 1 year, and 2 years. Shown are total brain volume and volumes of the olfactory bulb, frontal part of cortex, hippocampus, and cerebellum. Each point represents an individual animal; bars indicate mean ± SD. Fold differences between wild type and *Ttll1*^-/-^ are indicated. Statistical analyses were performed by two-tailed unpaired *t*-tests; *p* values are indicated. **d)** Longitudinal analysis of lateral ventricle volume in wild-type and *Ttll1*^-/-^ mice measured by MRI at 3, 6, 10, and 18 months. Each point represents an individual animal; bars indicate mean ± SD. Fold differences between wild type and *Ttll1*^-/-^ are indicated. Statistical analyses were performed by two-tailed unpaired *t*-tests; *p* values are indicated.

To quantify volumetric changes of distinct brain regions, we performed high-resolution T2-weighted 3D magnetic resonance imaging (MRI). The analysis was performed postmortem, and whole heads were analysed to preserve the brain structure inside the skull. Given the absence of visible morphological changes in *Ttll7^-/-^* brains, and the similarities between *Ttll1^-/-^* and *Ttll1^-/-^Ttll7^-/-^* brains (Fig. 2a; S2), we decided to focus on wild-type and *Ttll1^-/-^* brains. MRI scans acquired with a 17.2 T Bruker Biospec preclinical scanner were manually segmented into the olfactory bulb, frontal part of the cortex, hippocampus, and cerebellum, followed by semi-automated quantification of their volumes with Fiji (Fig. 2b,c; S3; Movie S1). No morphological differences between wild-type and *Ttll1^-/-^* brains were found in 2-month-old brains (Fig. 2b,c; S3; Movie S1), whereas a clear reduction in the volume of the olfactory bulb and the frontal part of the cortex was seen in 1- and 2-year-old brains of *Ttll1^-/-^* mice (Fig. 2b,c; S3; Movie S1). The olfactory bulb volume of *Ttll1^-/-^* mice was ∼2-fold reduced, while the volume of the frontal part of the cortex was reduced ∼1.5 fold (Fig. 2c). By contrast, hippocampus and cerebellum merely showed some variability in *Ttll1^-/-^*mice, however without displaying a clear trend (Fig. 2c). We also measured the volume of lateral ventricles by MRI of living animals, as increased ventricular volume is indicative of cerebrospinal fluid (CSF) accumulation. We found a progressive enlargement in the volume of the lateral ventricles with age in *Ttll1^-/-^* mice (Fig. 2d), a known hallmark of neurodegenerative disorders^64, 65^.

The observed phenotypes, atrophy of some of the brain regions and concomitant enlargement of ventricles, together with their progressive occurrence, strongly imply the presence of late-onset neuropathology – most likely neurodegeneration - in mice lacking TTLL1. By contrast, knockout of TTLL7 does not cause visible brain morphology abnormalities, nor does it aggravate *Ttll1^-/-^*-linked phenotypes in the *Ttll1^-/-^Ttll7^-/-^* double-knockout context. This correlates with our earlier findings that abnormal accumulation of TTLL1-catalysed polyglutamylation in mice lacking the deglutamylase CCP1 is causative for the massive neurodegeneration in these mice, while TTLL7 had no impact on this type of degeneration^41^. It might imply that TTLL1 regulates key physiological processes, likely by controlling neuronal cargo transport^29, 37, 41^ and the binding of MAPs such as Tau^49^ or other neuronal MAPs^48^, which, if perturbed, results in the degeneration of the affected neurons.

### Reduced glutamylation causes late-onset loss of mature neurons

As knockout of *Ttll1* caused a strong reduction of the volume of the olfactory bulb and the frontal part of the cortex in aging mice, we set out to investigate the degenerative phenotype at the cellular level. We performed immunohistological analyses of brain sections of young (2-month-old) and aged (1- and 2-year-old) wild-type, *Ttll1^-/-^*, *Ttll7^-/-^*, and *Ttll1^-/-^Ttll7^-/-^* mice. First, we analysed the olfactory bulb using the NeuN antibody, a marker of mature neurons^66^. We found an age-dependent loss of NeuN-positive neurons in the glomerular layer of the olfactory bulb of *Ttll1^-/-^* mice (Fig. 3a; S4). In contrast to olfactory granule cells, which are replenished throughout life by the rostral migratory stream (RMS)^67^, at least some of the glomerular neurons are not replaced in the adult^68, 69^; thus, their loss, while not alone accounting for the massive volume reduction of the olfactory bulb, is an indication for neurodegeneration. Quantification of the number of NeuN-positive neurons in the glomerular layer confirmed a drastic loss of these cells in 1- and 2-year-old *Ttll1^-/-^* and *Ttll1^-/-^Ttll7^-/-^* mice (Fig. 3b), while their number remained at wild-type levels in *Ttll7^-/-^* mice. These results suggest that TTLL1, but not TTLL7, is essential for maintaining mature neurons in olfactory bulb glomeruli, and lack of TTLL1 leads to their progressive loss.

**Figure 3:**
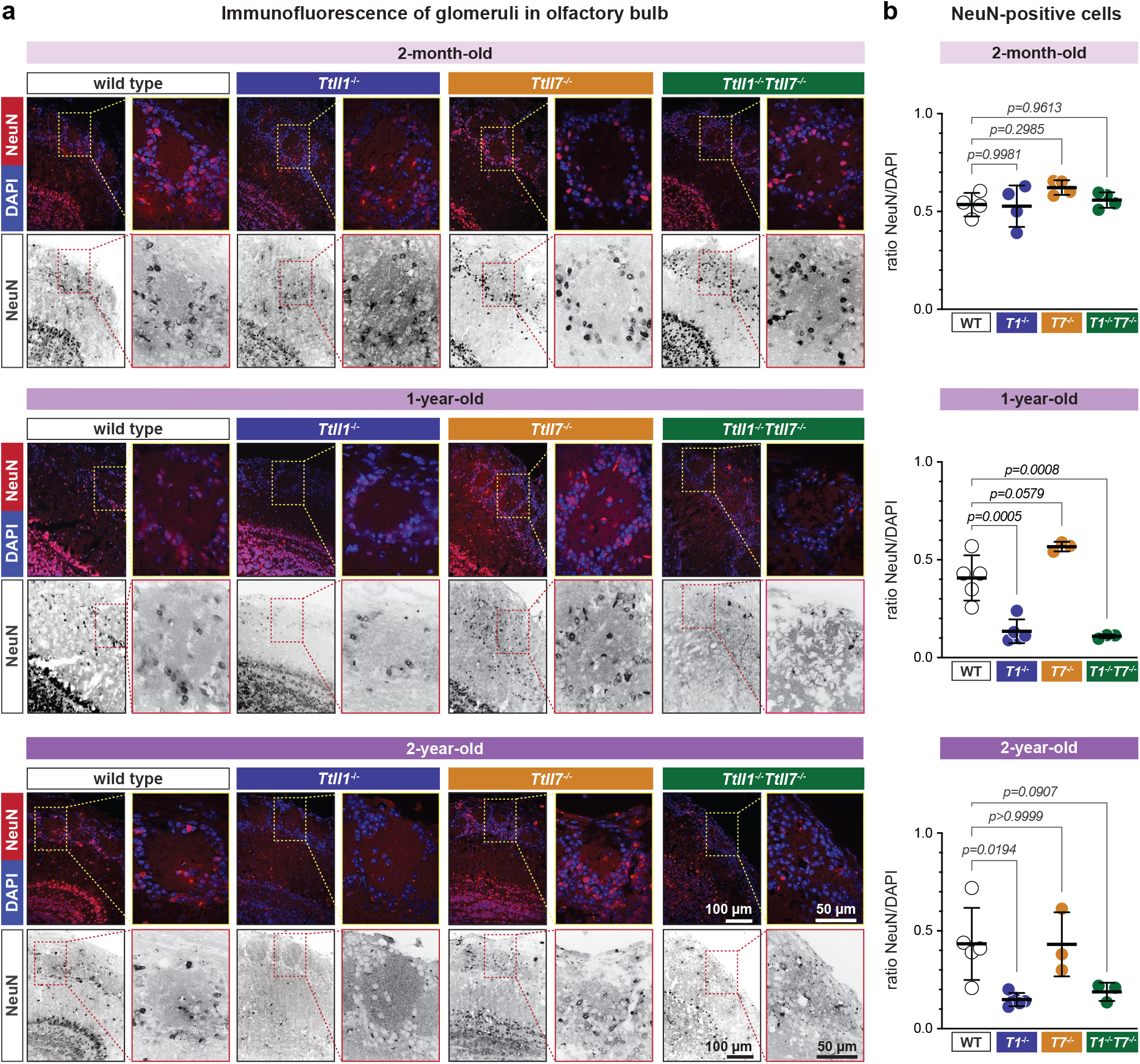
Immunohistological analyses of the olfactory bulb of mouse brains. **a)** Representative confocal images of NeuN immunostaining in the glomeruli of the olfactory bulb of wild-type, *Ttll1*^-/-^, *Ttll7*^-/-^, and *Ttll1*^-/-^*Ttll7*^-/-^ mice at 2 months, 1 year, and 2 years. NeuN (red) labels nuclei of mature neurons, DAPI (blue) marks all nuclei, and greyscale panels show NeuN single-channel images. For each animal, 20x (overview) and 40x (inset; from Fig. S4) images are shown. Scale bars: 100 μm (overview), 50 μm (inset). **b)** Quantification of the ratio of NeuN-positive cells vs. DAPI staining in olfactory-bulb glomeruli of wild-type (open circles), *Ttll1*^-/-^ (blue), *Ttll7*^-/-^ (orange), and *Ttll1*^-/-^*Ttll7*^-/-^ (green) for 2-month-old, 1-year-old, and 2-year-old mice. Each point represents an individual mouse; bars represent mean ± SD. Statistical analyses were performed by one-way ANOVA with multiple comparisons, *p* values are indicated.

We next explored the effect of reduced polyglutamylation in the frontal part of the cortex, which was also atrophied in aged mice lacking TTLL1 (Fig. 2). We determined the overall cell density in six different regions of the cortical sections by counting the number of DAPI-positive nuclei. No difference in the cell density could be found in wild-type, *Ttll1^-/-^, Ttll7^-/-^* and *Ttll1^-/-^Ttll7^-/-^* brains at all analysed ages (Fig. 4a,b; S5a). Analyses of mature neurons with NeuN staining further showed that the density of neurons was also unchanged across genotypes or ages (Fig. 4c,d; S5b). The same result was obtained when quantifying parvalbumin-positive interneurons (Fig. 4e,f; S6a). The constant density of neurons together with the concomitant decrease of brain volume clearly reveals an overall loss of neurons in mice lacking TTLL1.

**Figure 4:**
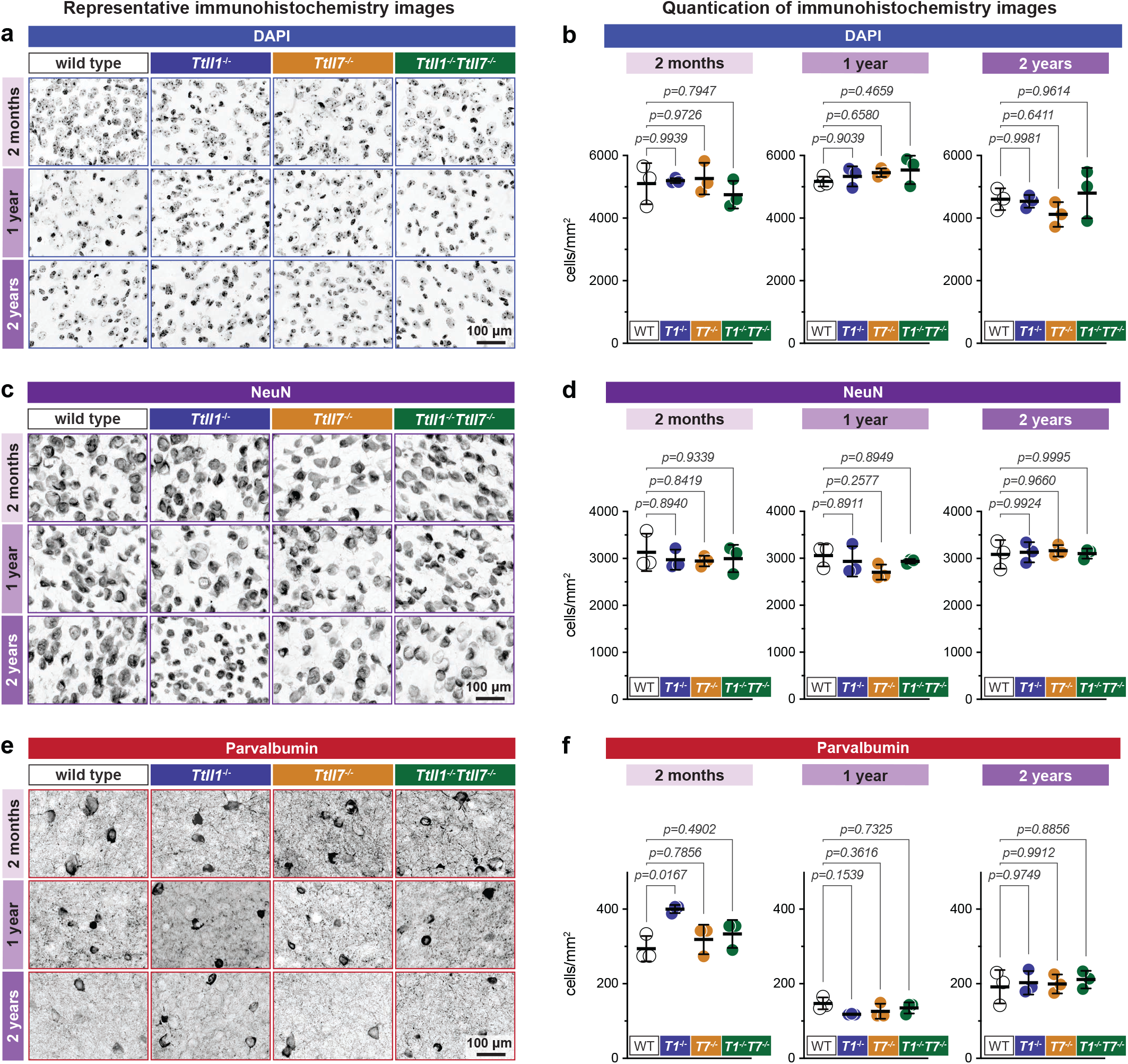
Immunohistological analyses of the frontal part of the mouse brain cortex. Different markers were used to estimate cell densities (in cells/mm^2^) in six representative regions of the frontal cortex part of wild-type, *Ttll1*^-/-^, *Ttll7*^-/-^, and *Ttll1*^-/-^*Ttll7*^-/-^ brains at 2 months, 1 year, and 2 years. **a, b)** Overall cell density. Representative DAPI staining (**a**) and quantification of total nuclei density (**b**). **c, d)** Density of mature neurons. Representative NeuN immunostaining (**c**) and quantification of neuronal density (**d**). **e, f)** Density of parvalbumin-expressing neurons. Representative Pvn immunostaining (**e**) and quantification of parvalbumin-positive interneurons (**f**). **a, c, e)** Images from Fig. S5. Scale bars: 100 μm. **b, d, f)** Wild-type animals are shown as open circles, *Ttll1*^-/-^ in blue, *Ttll7*^-/-^ in orange, and *Ttll1*^-/-^*Ttll7*^-/-^ in green. Each dot represents an individual animal; bars represent mean ± SD. Statistical analyses were performed by one-way ANOVA with multiple comparisons, *p* values are indicated.

### Reduced glutamylation causes an age-onset reduction of dendritic spine density

Given the late-onset and relatively moderate loss of neurons in the frontal cortex, we aimed at analysing earlier markers of neuronal defects. We investigated synaptic densities, given that synapse loss is known to be associated with cortical atrophy^70^. We used DiOlistic labelling^71, 72^, a technique in which single neurons in fixed thick brain sections are filled with a lipophilic fluorescent dye to visualise their morphological details. We used this technique to quantify dendritic spine densities of pyramidal neurons in the frontal cortex and hippocampus of wild-type and *Ttll1^-/-^* mouse brains at 2 months and 1 year.

In the frontal cortex of 2-month-old mice, we found a slight increase of ∼10% in spine number (Fig. 5a,b; S6a,c). This is in accordance with previous findings of increased spine density in hippocampal neurons of *Ttll1^-/-^* mice of this age linked to delayed pruning^73^. By contrast, 1-year-old *Ttll1^-/-^* animals showed a reduction of about ∼16% in spine density in the frontal cortex (Fig. 5a,b; S6b,d), which correlates with the atrophy observed in this part of the brain. To assess whether synaptic defects extended beyond the frontal cortex, we also counted dendritic spines in the pyramidal neurons of the hippocampal dentate gyrus in 2-month-old and 1-year-old mice. Similar to cortical neurons, hippocampal neurons from *Ttll1^-/-^* brains displayed an increased spine density of ∼11% at 2 months (Fig. 5c,d; S7a,c), while at 1 year, spines were reduced by ∼20% (Fig. 5c,d; S7b,d), highlighting an age-dependent synaptic vulnerability caused by TTLL1 depletion and reduced tubulin polyglutamylation.

**Figure 5:**
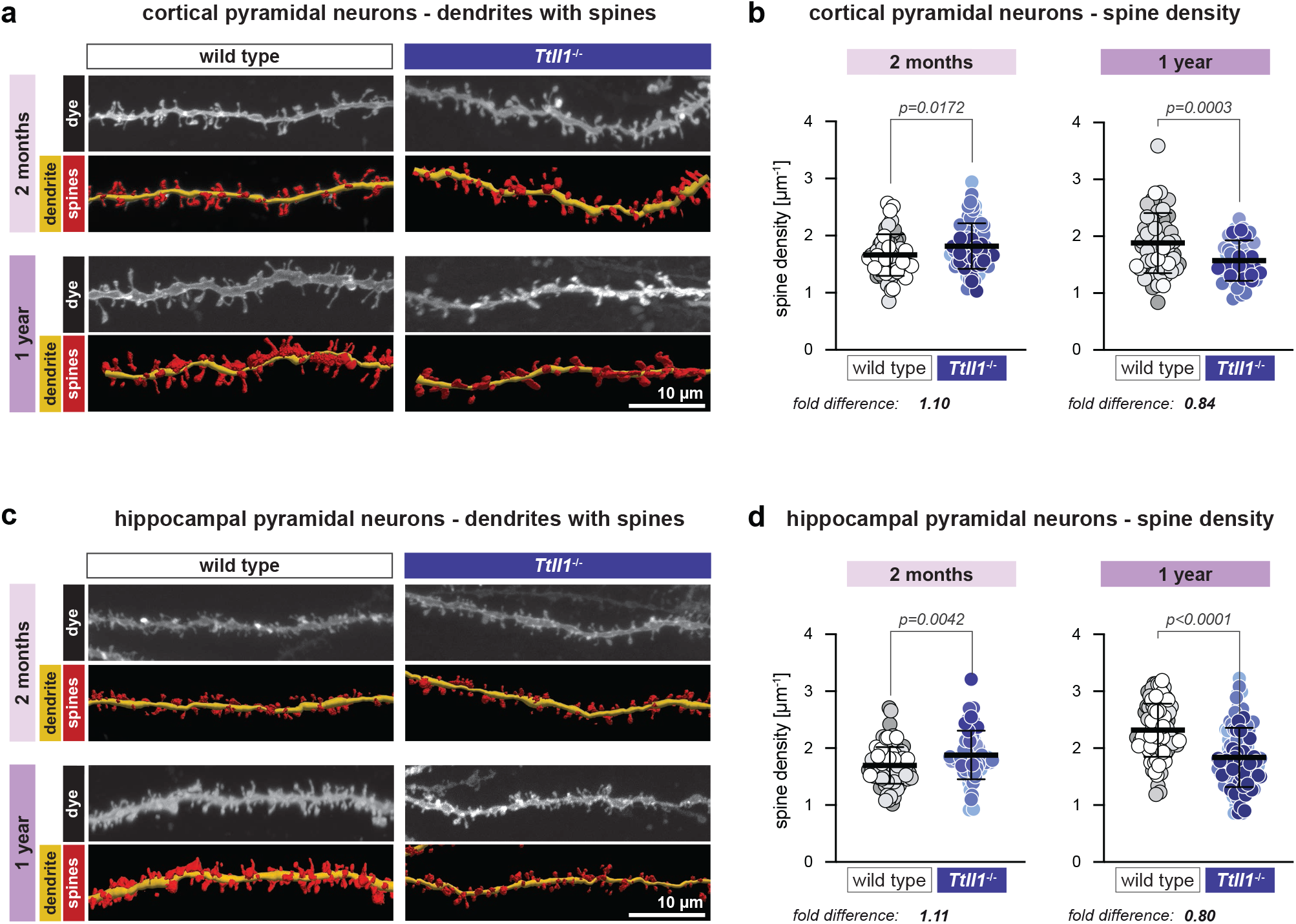
Impact of reduced polyglutamylation on spine density. **a)** Representative images of dendritic segments from cortical pyramidal neurons of wild-type and *Ttll1*^-/-^ mice at 2 months and 1 year (images for single analysed animals are shown in Fig. S6a,b). Upper panels show greyscale images of dye-filled dendrites; lower panels show 3D reconstructions with dendrites in yellow and dendritic spines in red. Scale bar: 10 μm. **b)** Quantification of dendritic spine density (spines/μm) shown in (**a**). Each dot represents one analysed dendrite. Results obtained from different animals are shown in different colour shades (white/grey: wild type; blue: *Ttll1*^-/-^; corresponding data from single animals are shown in Fig. S6c). Bars represent mean ± SD; fold differences are indicated. Statistical analyses were performed with unpaired *t*-test with Welch’s correction, *p* values are indicated. **c)** Representative images of dendritic segments from hippocampal pyramidal neurons of wild-type and *Ttll1*^-/-^ mice at 2 months and 1 year (images for single analysed animals are shown in Fig. S7a,b). Upper panels show greyscale images of dye-filled dendrites; lower panels show 3D reconstructions with dendrites in yellow and dendritic spines in red. Scale bar: 10 μm. **d)** Quantification of dendritic spine density (spines/μm) shown in (**c**). Each dot represents one analysed dendrite. Results obtained from different animals are shown in different colour shades (white/grey: wild type; blue: *Ttll1*^-/-^; corresponding data from single animals are shown in Fig. S7c). Bars represent mean ± SD; fold differences are indicated. Statistical analyses were performed with unpaired *t*-test with Welch’s correction, *p* values are indicated.

Together, these findings demonstrate that absence of TTLL1-generated polyglutamylation induces an age-onset loss of synapses, a sign of perturbed neuronal homeostasis in the frontal cortex, but also in a region that did not show obvious signs of degeneration, the hippocampus.

### Reduced polyglutamylation affects behavioural and cognitive functions

Neuropathology is often accompanied by the perturbation of neuronal circuits, which can result in behavioural and cognitive abnormalities^74^. Furthermore, cognitive impairments can be associated to the malfunction of neurons before the cellular effects can be detected. To investigate the impact of reduced polyglutamylation at the behavioural level, we evaluated olfaction as well as learning and memory of our mouse models.

We first evaluated the functional consequences of olfactory bulb degeneration caused by *Ttll1* deletion using an olfaction assay^75^. The test assesses the reaction of animals to two non-social odours (almond and banana extracts) and two social odours (urine-derived pheromones from same- and neighbouring-cage mice). At 2 months, mice of all genotypes were capable of detecting all odours (Fig. S8a). At 1 year, *Ttll1^-/-^* and *Ttll1^-/-^Ttll7^-/-^* mice exhibited deficits in detecting both social and non-social odours, which was not the case for wild-type and *Ttll7^-/-^*mice (Fig. S8a). This, together with the observed loss of mature neurons in the olfactory-bulb glomeruli of animals lacking TTLL1 (Fig. 3; S4), suggests that TTLL1-mediated polyglutamylation is indispensable for maintaining olfaction in mice.

To explore cognitive functions, we used the automated IntelliCage^®^ system^76^ (Fig. S8b), which, in contrast to classical behavioural tests, allows testing mice in their social group with minimal interference from the experimenter. Wild-type, *Ttll1^-/-^* and *Ttll7^-/-^* mice of 2 months, 1 and 2 years were introduced in the IntelliCage^®^ (Fig. S8b) to evaluate the different aspects of their cognitive behaviours such as learning, long-term and spatial, memory, and impulsivity (Fig. 6a, S8). Up to 12 animals of the same sex and age, but different genotypes, were tested together in each IntelliCage^®^. During the first days of housing in the IntelliCage^®^, while the animals were adapting to the novel environment, we assessed spontaneous and exploratory behaviour by measuring the daily average number of total corner visits (Fig. S8c) and the daily average number of visits without nose pokes (Fig. 6b), respectively. Young *Ttll1^-/-^* mice showed a reduced number of corner visits and visits without nose poke as compared to wild-type mice (Fig. 6b, S8c), suggesting a reduced basal activity and an increased anxiety-like trait.

**Figure 6:**
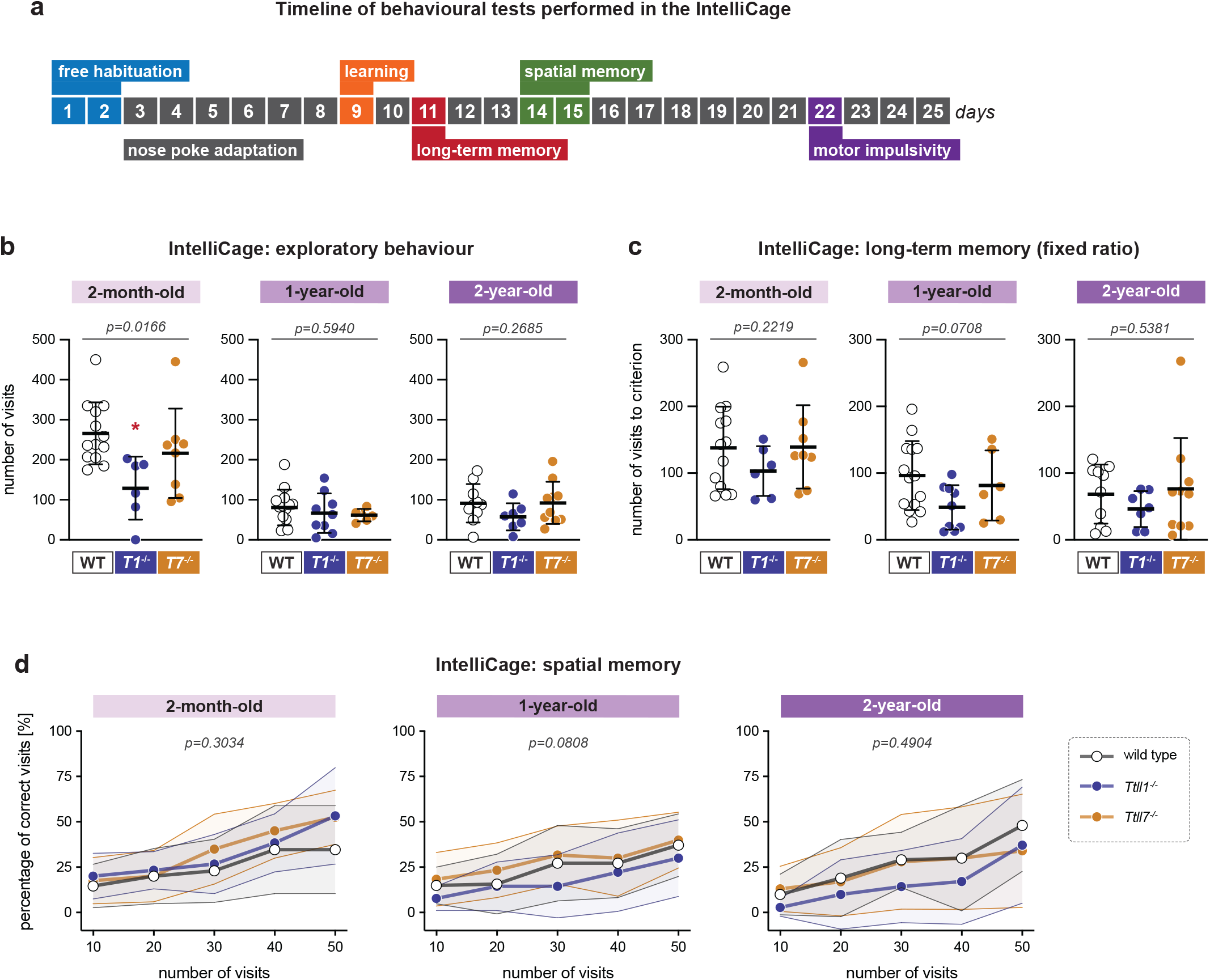
Behavioural effects of reduced polyglutamylation. **a)** Schematic representation of the sequence and duration of the different tests performed in the IntelliCage^®^ shown in (**b, c, d**) and Fig. S8c,d,e. **b, c, d)** Behavioural assessment carried out in the IntelliCage^®^ on 2-month-old, 1- and 2- year-old wild-type, *Ttll1*^-/-^ and *Ttll7*^-/-^ mice. Each dot represents one animal; bars represent mean ± SD. **b)** Quantification of exploratory behaviour. Number of visits without nose pokes to the corners of the cage is recorded during the first 3 days in the IntelliCage^®^. Higher values indicate elevated exploratory activity. Data were analysed using one-way ANOVA followed by Tukey’s post hoc test. *P* value indicates overall statistical significance of differences across genotypes. * p<0.05 *Ttll1^-/-^*vs. wild-type. **c)** Results of the fixed ratio paradigm as assessment of the long-term memory. Number of visits performed to obtain water access is plotted. Lower values indicate better long-term memory. Data were analysed using one-way ANOVA. *P* value indicates overall statistical significance of differences across genotypes. **d)** Evaluation of spatial learning capabilities. The percentage of visits to the correct corner is plotted. Higher values indicate better spatial learning. Data were analysed using two-way ANOVA. *P* value indicates overall statistical significance of differences across genotypes.

Following one day of progressive-ratio training (Fig. S8d), long-term memory was evaluated using a fixed-ratio schedule (Fig. 6c). The number of visits required for each animal to reach the nose pokes requirement achieved during the previous day’s training was used as an index of procedural memory retention. Across all ages analysed, *Ttll1^-/-^* mice consistently tended to reach the breakpoint earlier than the wild-type and *Ttll7^-/-^* mice (Fig. 6c). To verify whether this behaviour was linked to an increased impulsivity of *Ttll1^-/-^* mice, we tested motor impulsivity by assessing the capacity to refrain from nose poking for water access for increasing time delays. Overall, there were no significant changes in impulsivity of the different mouse strains, indicating that this behaviour is not affected by the genotype (Fig. S8e). While the results of the fixed-ratio test (Fig. 6c) were not statistically significant, they suggest, together with the absence of increased impulsivity (Fig. S8e), that *Ttll1^-/-^* mice might have better long-term memory.

We next analysed spatial memory (Fig. 6d). In this test, access to water was restricted to one specific corner of the cage and the percentage of visits to the correct corner was used to evaluate spatial memory. While 2-month- and 1-year-old *Ttll1^-/-^*mice showed behaviour similar to the wild-type and *Ttll7^-/-^* mice, at 2 years their performance was less efficient (Fig. 6d), suggesting a progressive decline in spatial memory.

Overall, the behavioural experiments demonstrate that *Ttll1^-/-^*but not *Ttll7^-/-^* mice display cognitive and behavioural alterations across their lifespan. Young *Ttll1^-/-^* mice showed reduced spontaneous and exploratory activity, indicative of altered adaptation to a novel environment. *Ttll1^-/-^* mice exhibited a consistent trend towards faster acquisition of the breakpoint in the long-term memory task across ages, whereas aged animals showed reduced spatial memory performance. These findings highlight the importance of TTLL1-mediated polyglutamylation for maintaining normal cognitive functions throughout life.

### Lack of TTLL1-mediated polyglutamylation leads to reduced survival of mice transgenic for human mutant P301S Tau

Polyglutamylation has been shown to control the binding of the neuronal MAP Tau to microtubules^49^. Pathological changes of Tau are hallmarks of a family of neurodegenerative disorders known as tauopathies, which famously include Alzheimer’s disease^58^. While the majority of spontaneous tauopathy cases are not linked to mutations, familial cases of these disorders carry disease-causing mutations^77^. One of them, the P301S amino acid substitution in Tau, is linked to familial frontotemporal dementia (FTDP-17)^78^. A murine model overexpressing human Tau^P301S^ develops severe neurodegeneration, including formation of filamentous Tau lesions, severe paralysis around 6-7 months, and reduced overall lifespan of less than 9 months^59^. The *Tau^P301S^* mouse can thus be considered as an accelerated tauopathy model, allowing us to test mechanisms linked to late-onset tauopathies within an experimentally accessible time window. We used this mouse model to determine the impact of polyglutamylation on the progression of tauopathies.

We generated a combinatorial mouse line *Tau^P301S^Ttll1^-/-^* and tested the primary readout of the degenerative phenotype – animals’ lifespan (Fig. 7a). Strikingly, while a cohort of 38 *Tau^P301S^* mice had a median survival of 197 days, a cohort of 20 *Tau^P301S^Ttll1^-/-^* showed a median survival of only 170 days (Fig. 7a), representing an almost 10% reduction of lifespan. These results demonstrate a potential mechanistic link between Tau pathology and polyglutamylation in an established model of tauopathy.

**Figure 7:**
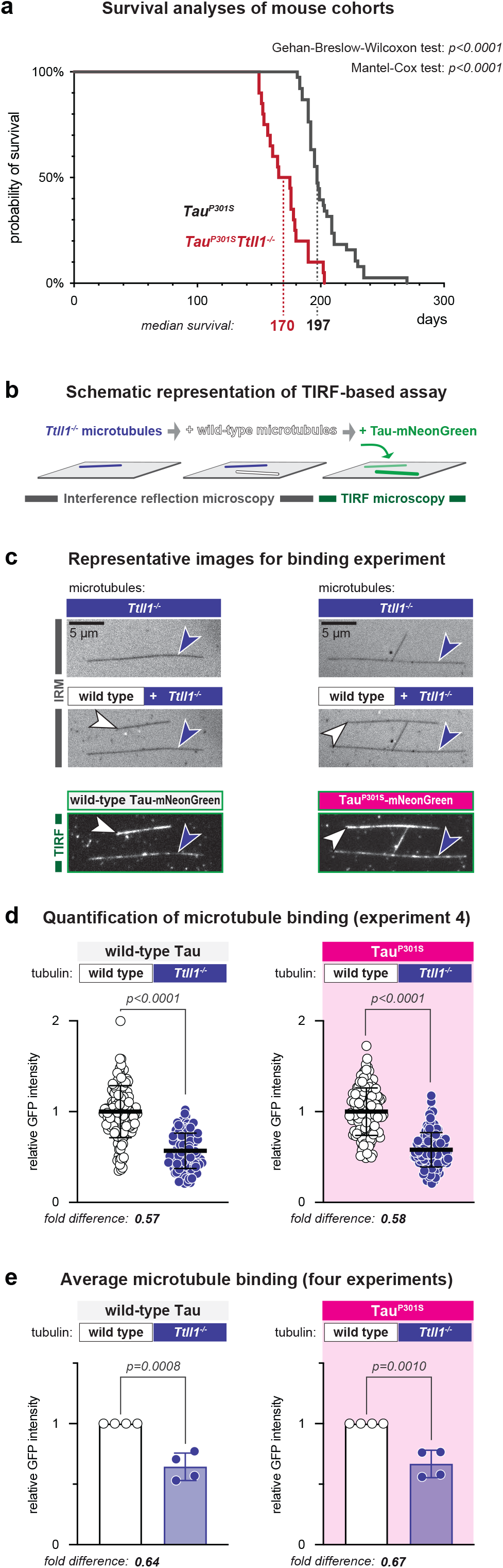
Reduced polylutamylation and Tau pathology. **a)** Kaplan-Meier curves showing the probability of survival of *Tau^P301S^* (grey) and *Tau^P301S^Ttll1^-/-^*(red) mice. Median survival of the two strains is shown below. Data were analysed using the Gehan-Breslow-Wilcoxon and the Mantel-Cox tests, *p* values are shown. **b-e)**Total Internal Reflection Fluorescence (TIRF) microscopy-based i*n vitro* analysis of the binding of mNeonGreen-tagged wild-type Tau and its mutated variant Tau^P301S^ to microtubules from either wild-type or *Ttll1^-/-^* brains. **b)**Schematic representation of experimental pipeline: microtubules are polymerised from tubulin purified from wild-type or *Ttll1^-/-^* brains, sequentially attached to a cover slip and imaged by interference reflection microscopy (IRM). Subsequently mNeonGreen-tagged Tau is added and fluorescent intensities on both types of microtubules are quantified. **c)**Representative microscopy images one experiment as described in (**b**). **d)**Quantification of wild-type Tau and Tau^P301S^ binding intensities to wild-type (open circles) and *Ttll1^-/-^* (blue); one experiment out of four replicates shown in Fig. S9, with fluorescence values normalised to the mean value of wild-type microtubules. Each data point represents a single microtubule. Bars represent mean ± SD. Fold differences are indicated. Statistical analyses were performed by two-tailed unpaired *t*-tests; *p* values are indicated. **e)**Average wild-type Tau and Tau^P301S^ binding to different microtubule types. Each data point represents the mean value from one of four independent experiments shown in Fig. S9. Fold differences are indicated. Statistical analyses were performed by two-tailed unpaired *t*-tests; *p* values are indicated.

To explore the mechanistic basis of this effect, we characterised the direct binding of Tau^P301S^ to microtubules using a Total Internal Reflection Fluorescence (TIRF) microscopy-based *in vitro* reconstitution approach (Fig. 7b). Purified mNeonGreen-tagged wild-type and Tau^P301S^ proteins were tested for their interactions with microtubules assembled from either wild-type or *Ttll1^-/-^* brain tubulin^79^. The two types of microtubules were immobilised in the same microscopy chamber, and the microtubule-binding of mNeonGreen-tagged Tau was directly measured and quantified on single microtubules (Fig. 7b,c)^49^. Four independent sets of experiments (Fig. 7d; S9) consistently showed (Fig. 7e) that both wild-type Tau and Tau^P301S^ interact more strongly with wild-type brain microtubules that are normally polyglutamylated (Fig. 1d). By contrast, their interactions with *Ttll1^-/-^*microtubules, which have reduced polyglutamylation on α-tubulin (Fig. 1d), were reduced by more than 30% (Fig. 7e). These results demonstrate that microtubule polyglutamylation controls the binding of the pathological Tau variant similar to wild-type Tau. Given that Tau aggregation is the main driving factor of neurodegeneration induced by Tau^P301S^, our findings imply that the reduced lifespan of *Tau^P301S^Ttll1^-/-^* mice, which suggests a more rapid time course of degeneration, is linked to reduced Tau-microtubule interactions. Polyglutamylation could therefore be a modulating factor in Tau-induced neurodegeneration.

## Discussion

The microtubule cytoskeleton is essential for development, homeostasis, and long-term survival of neurons. Consequently, defects of microtubules and their interacting proteins are associated with both neurodevelopmental and neurodegenerative diseases. While in many neurodegenerative disorders the described culprits are dysfunctional microtubule interactors, such as molecular motors^4, 8, 80^, structural MAPs^56, 58^, or microtubule-severing enzymes^54^, the role of microtubules themselves has only recently been addressed. Tubulin PTMs, which have long been suspected to regulate microtubule functions, emerged as potential players in neuropathologies. In particular polyglutamylation, abnormal accumulation of which causes early-onset neurodegeneration in mice and humans^29, 42^, can be considered a promising candidate. Polyglutamylation is enriched in differentiated neurons^35, 36^ and the expression of modifying enzymes is lower in development as compared to adult neurons and brain^37^. This is different for another tubulin PTM, tubulin detyrosination, which shows highest enzyme expression levels during neuronal development^37^ and perturbation of which leads to severe developmental defects^26^, but which does not cause neurodegeneration when perturbed later in development^39, 81^.

Physiological levels of polyglutamylation in neurons are relatively high – most of the tubulin molecules isolated from brain are modified^32–34, 63^, suggesting that this PTM is required for neuronal microtubule functions. This together with the finding that abnormal accumulation of polyglutamylation causes massive neurodegeneration^29, 42^ suggested that removal of this PTM should reveal its role in neuronal function and survival. We analysed mice lacking two polyglutamylases, which we had shown generate most of the polyglutamylation found in the brain, TTLL1 and TTLL7; TTLL1 polyglutamylating α- and TTLL7 β-tubulin^41, 49^. Mice lacking both enzymes show very low overall levels of polyglutamylation in the brain, suggesting the absence of compensation between glutamylating enzymes. Importantly, knockout of *Ttll1* and *Ttll7* did not substantially alter other tubulin PTMs, such as acetylation or detyrosination, highlighting the independent regulation of different tubulin PTMs at the organism level.

Longitudinal analysis of our knockout mouse models revealed that certain parts of the brains of *Ttll1^-/-^* and *Ttll1^-/-^Ttll7^-/-^*, but not of *Ttll7^-/-^* animals, underwent age-dependent atrophy. This indicated that polyglutamylation generated by TTLL1 has a more prominent role for healthy neuronal ageing than that generated by TTLL7. These observations mirrored the predominant role of TTLL1 in the abnormal accumulation of polyglutamylation in mice lacking CCP1, which in turn causes the severe neurodegeneration in these mice^41^. While mechanisms leading to the pathology in mice with increased or decreased polyglutamylation might differ, in both cases we found alterations in axonal transport, a well-recognised hallmark of neurodegenerative disorders^4, 8^. Again, TTLL1 was the polyglutamylase associated with transport perturbations: reduced transport in hyper-^29^ and enhanced transport in hypo-glutamylation^41^.

*Ttll1^-/-^* mice developed late-onset neuropathology highly reminiscent of degeneration in different regions of the nervous system than those affected in *Ccp1^-/-^* mice, which is unsurprising given that even related neurodegenerative disorders can have different phenotypical appearances^82, 83^. In contrast to the early-onset degeneration in *Ccp1^-/-^* mice, *Ttll1^-/-^*brains appeared normal at 2 months but showed atrophy coupled to neuronal loss in the olfactory bulb and frontal parts of the cortex in 1-year-old animals. We demonstrated that mature rather than developing neurons were lost in the olfactory bulb, which underpins that the observed neuronal loss is likely to be caused by neurodegeneration. This does not exclude that perturbed migration of neuronal progenitors in the rostral migratory stream that replenish neurons in this brain region also contributes to the observed volume loss of the olfactory bulb.

The presence of a rather subtle neuropathology could make *Ttll1^-/-^*mice a valid model for late-onset neurodegeneration. This possibility was further underpinned by the synaptic loss we found in 1-year-old *Ttll1^-/-^* mice, which is considered an early event in neurodegeneration^84, 85^. Strikingly, young animals showed increased numbers of spines in *Ttll1^-/-^*mice, which, as described previously, is due to delayed pruning^73^. This mirrors that polyglutamylation, by controlling multiple microtubule-based processes, can have opposing impacts on neuronal connectivity depending on the age of the animal. During developmental pruning, polyglutamylation is important for controlling the activity of the microtubule-severing enzyme spastin^73^, hence the delayed pruning in the absence of TTLL1. By contrast, reduction of synapse numbers in ageing neurons could be caused by the previously demonstrated perturbed axonal transport in *Ttll1^-/-^* neurons^41^. Controlled axonal transport of synaptic vesicles is indeed essential for the proper function and maintenance of synapses^86^. At the same time, reduced interactions of Tau with microtubules^49^ could play a part in polyglutamylation-induced synaptic loss, given that pathological Tau^P301S^ causes a similar phenotype^87^.

We further explored cognitive phenotypes that are often observed before cellular changes are visible in the brain^88^. Cognitive assessments revealed a complex phenotype in *Ttll1^-/-^* mice, including reduced exploratory activity already in 2-month-old animals, preserved long-term memory, but deficits in spatial memory in 2-year-old animals. This suggested that Ttll1-mediated polyglutamylation is important for maintaining hippocampal and prefrontal circuit function underlying memory and cognitive control. Our tests did not show notable behavioural perturbations in the *Ttll7^-/-^* mice, which aligns with the absence of morphological abnormalities in these mice.

Having established reduced polyglutamylation as a cause of age-onset neuropathology, a final question was whether this PTM can directly contribute to neurodegeneration, specifically tauopathy, the most frequent cause of late-onset degeneration in humans. We tested this by combining a P301S Tau mutation, linked to familial cases of frontotemporal dementia^78^, with *Ttll1^-/-^*. *Tau^P301S^* mice show many hallmarks of tauopathy such as formation of neurofibrillary tangles, and most remarkably, early demise^59, 89^. Moreover, Tau^P301S^ triggers loss of synaptic Tau accompanied by decreased dendritic spine density^87, 89, 90^ similar to our findings in *Ttll1^-/-^* mice. Our observation that the genetic combination of P301S-mutated Tau with reduced polyglutamylation further shortens the life span of these mice suggests that Tau pathology becomes more severe when the Tau-microtubule interactions are weakened on microtubules lacking TTLL1-mediated polyglutamylation. Interestingly, another FTD-causing Tau mutant mouse model (*MAPT^S305N;Int10+3G>A^*) shows similar cellular and behavioural phenotypes to the *Ttll1^-/-^*mouse without the presence of neurofibrillary tangles^91^. This suggests that damage inflicted on neurons by progressing Tau pathology can be detrimental at the cellular and organism level before the apparition of aggregated Tau or neurofibrillary tangles^89, 92^. It is thus possible that decreased polyglutamylation levels, by reducing Tau binding to microtubules, can cause neuronal dysfunction even in the absence of mutated Tau, but not in the time windows observable within the lifespan of mice. Our results also align with a previous study in which polyglutamylation was reduced by site-directed mutagenesis rendering the α-tubulin isotype TUBA4A impossible to polyglutamylate by TTLLs^93^. The decrease of α-tubulin polyglutamylation in this mouse model was less prominent than in the here-studied mouse lacking TTLL1, which is unsurprising given that TUBA4A represents only about one third of α-tubulin isotypes expressed in the adult brain^94^. Nonetheless the authors observed several signs of neuropathology underpinning the importance of α-tubulin polyglutamylation for neuronal homeostasis^93^.

In conclusion, our characterisation of mouse models lacking polyglutamylation in the ageing brain revealed an essential function of this PTM for neuronal homeostasis. Absence of Ttll1-generated polyglutamylation engendered behavioural deficits in these animals and led to progressive loss of dendritic spines, both of which are early signs of neuronal dysfunction.

The presence of a neuropathology was ultimately revealed by a progressive loss of neurons in the olfactory bulb and the frontal cortex. We further demonstrate that decreased polyglutamylation aggravates the progression of the pathology in an established mouse model for tauopathy. Our work thus demonstrates that reduced polyglutamylation is a potential pathogenic factor in the development of neurodegenerative disorders. The fact that it affects a fundamental molecular process, the control of microtubule interactions with molecular effectors, might make it therapeutically exploitable for corrective interventions at early stages of neurodegenerative disorders such as tauopathies that affect a significant proportion of the ageing human population.

## Supporting information

Chakraborty MovieS1

## Acknowledgments

This work was supported by the Institut Curie, the Centre National de la Recherche Scientifique, the Agence Nationale de la Recherche ANR-10-IDEX-0001-02 and the LabEx Cell(n)Scale ANR-11-LBX-0038 and the European Research Council (ERC-2022-SYG) under the European Union’s Horizon 2020 research and innovation programme (grant agreement N°101071583 ‘TUBULINCODE’ to CJ, ZL). CJ and IB are supported the French National Research Agency (ANR) award ANR-20-CE13-0011, CJ by the Fondation pour la Recherche Medicale (FRM) grants MND202003011485 and EQU202203014694. MMM is supported by the Fondation Vaincre Alzheimer FR-16055p and the France Alzheimer grant 2023. SC was supported by a PSL-Biogen PhD grant via a CIFRE contract. We further acknowledge institutional support from the CAS (RVO: 86652036), Centre of Molecular Structure supported by MEYS CR (LM2015043), the Imaging Methods Core Facility at BIOCEV supported by the MEYS CR (Large RI Project LM2018129 Czech-BioImaging), ERDF (project no. CZ.02.1.01/0.0/ 0.0/16_013/0001775) to ZL, support by the Czech Science Foundation grant 25-17813S and the IPHYS Bioimaging Facility supported by BIF MEYS CR (Large RI Project LM2023050 Czech-BioImaging) and ERDF (Project No. CZ.02.1.01/0.0/0.0/18_046/0016045) to MB.

We thank C. Alberti, E. Belloir, K. Belloul, V. Dangles-Marie, C. Jouhanneau, A. Luber (Institut Curie), as well as E.-A. Benlarbi, R. Gaudin and W. Nguyen (Neurospin, CEA, Gif-sur-Yvette) for technical assistance, as well as M.-N. Soler and C. Messaoudi from the Multimodal Imaging Center of the Institut Curie (CNRS UAR2016/ Inserm US43/ Institut Curie / Université Paris-Saclay) and S. Leboucher from the histology platform (Institut Curie, Orsay). We are grateful to C. Coarfa, C. Walker (Baylor College of Medicine, Houston USA), P. Bost, M. Corbe, F. Mangane and N. Servant (Institut Curie, Paris) for single-cell analyses. We thank H. Hering, D. Walsh, and M. Beyna (Biogen, Cambridge, USA) for their constant support of our project, for hosting SC in their team, and for teaching essential skills required for the current project.

We are grateful to A. Baffet (Institut Curie, Paris), M. Braun (IBT, CAS, Prague), N. Renier (Institut du Cerveau, Paris), F. del Bene (Institut de la Vision, Paris), Y. Song (Massachusetts General Hospital, Charlestown, USA), N. Mizuno (NIH, Bethesda USA), T.J. Hausrat (ZMNH, Hamburg, Germany), Y.-C. Lin (National Tsing Hua University, Hsinchu City, Taiwan), J. Howard (Yale University, New Haven, USA), C. Walker (Baylor College of Medicine, Houston, USA), C. Gomes da Silva (UMC, Utrecht, The Netherlands), G. Schiavo (UCL, London, UK) for the insightful discussion, suggestions and support in analysing and interpreting our data.

## Materials and Methods

### Mouse breeding

Animal care and use for the experiments were performed in accordance with the recommendations of the European Community (2010/63/UE). Experimental procedures were specifically approved by the ethics committee of the Institut Curie CEEA-IC #118 (authorisations APAFIS #04395.03, #37315-2022051117455434 v2 and #32940-2021091316059579 v1 given by National Authority) in compliance with the international guidelines. Young adult (2- and 3-months), adult (6-, 10-, 12-, and 17-months) and aged (24-months) females and males were used in this study.

Work involving mice at the MRC Laboratory of Molecular Biology was carried out under Home Office PPL PP8626039, with approval from the LMB’s Animal Welfare and Ethical Review Body.

### Mouse lines

C57BL/6NTac-*Ttll1^tm1a(EUCOMM)Wtsi^/IcsOrl* mice were generated at PHENOMIN-ICS (Institut Clinique de la Souris, Illkirch, France; www.ics-mci.fr) with an ES-cell clone obtained from the International Mouse Phenotyping Consortium (IMPC) (www.mousephenotype.org/data/alleles/MGI:2443047/tm1a(EUCOMM)Wtsi), and bred first to flp-, then to cre-recombinase-expressing animals^95^ to create the *Ttll1*^-/-^ line as described before^29^. The *Ttll7*^tm1Ics^ conditional mutant mouse line was established at PHENOMIN-ICS (Institut Clinique de la Souris, Illkirch, France; www.ics-mci.fr) as a conditional (Lox-P) line and then bred to PGK-cre-recombinase animals^95^ to create the *Ttll7*^-/-^ line as described before^41^.

### Mouse genotyping

For adult mice, DNA was extracted from ear fragments collected during the identification of mice or from tail fragments using lysis buffer (0.1 M Tris–HCl pH 8 (Sigma #T1503), 0.2 M NaCl (Sigma #S3014), 5 mM EDTA (Euromedex #EU0007-C) and 0.4% SDS (Euromedex #EU0660)) containing 0.1 mg/ml proteinase K (#193504, MP Biomedicals). Lysis buffer with proteinase K was added to the sample and incubated at 56°C for 4 h or overnight. After the digestion, the proteinase K was inactivated by boiling the sample at 95°C for 10 min

### Antibody generation

The sequence for the C-terminal tail peptide of β2-tubulin (TUBB2A) was designed with glutamylation on the amino acid residues that are found to be highly modified in brain tissue^33^ (Fig. 1b). A standard peptide synthesis protocol was used to generate peptides with a bi-glutamylated side chain on glutamate 435. The peptide was purified by high-performance liquid chromatography (HPLC; Peptide Specialty Laboratory, Heidelberg, Germany) and coupled to keyhole limpet hemocyanin, and polyclonal antibodies were raised in rabbits. For purification, 3 mg of the modified peptide was covalently linked to an NHS-activated Sepharose column (#GE17-0716-01; Sigma-Aldrich) as per the manufacturer’s protocol. The blood serum was centrifuged at 90,000×*g* for 30 min at 4°C to remove any cell debris, filtered with a 0.2-µm syringe filter (#16532; Sartorius) and loaded onto the affinity column. After washing with ∼20 ml of PBS, the antibodies were eluted with 100 mM glycine-HCl, pH 2.3, which was immediately neutralised to pH 8.0 with 1 M glycine, pH 9.0. The purified antibody was subsequently dialysed against 3×1 l PBS at 4°C. The concentration of the antibody was estimated with the Pierce BCA protein assay kit (#23225; Life Technologies).

### Sample preparation and immunoblotting

Mice of desired genotype and age were sacrificed by cervical elongation, and brain tissue was immediately dissected. Brain tissues from wild-type, *Ttll1*^-/-^, *Ttll7*^-/-^ and *Ttll1*^-/-^*Ttll7*^-/-^ mice were collected in 2.5× Laemmli buffer: 180 mM DTT (Sigma #D9779), 4% SDS (VWR #442444H), 160 mM Tris–HCl pH 6.8, 20% glycerol (VWR #24388.295), bromophenol blue, and homogenised using an Eppendorf tube potter. The samples were boiled at 95°C for 5 min, spun down at 15,000×*g* for 5 min using a tabletop centrifuge, and supernatants were stored at -20°C. Samples were run on SDS–PAGE gels, allowing to separate α- and β-tubulin^60^ and transferred onto a nitrocellulose membrane using a Bio-Rad Trans-Blot^®^ Turbo system, according to the manufacturer’s instructions. The membranes were blocked in 5% skimmed milk prepared in 1×TBST (Tris-buffered saline containing 0.1% Tween 20 (VWR #0777)) and incubated with desired primary antibodies for 2 h (see Appendix Table S1 for dilutions). Chemiluminescence signal on the membrane was revealed using Clarity™ Western ECL Substrate (Bio-Rad #1705060) solution and the Vilber imaging system.

### Transcardial perfusion

Animals were anaesthetised using ketamine (Imalgène^®^, 80–100 mg/ml)/xylazine (Rompun^®^, 5–10 mg/ml) and transcardially perfused first with PBS followed by 4% PFA (diluted from 32% EM grade, #15714, EMS) in phosphate buffer (PB; 100 mM Na_2_HPO_4_/NaH_2_PO_4_ pH 7.4) and post-fixed in 10% neutral buffered formalin (Qpath, #FOR0020AF59001) for 48 h at room temperature for immunohistochemistry. For MRI imaging, mice were perfused with 30 ml of a fixative solution containing 2% PFA, 2% glutaraldehyde, and 120 µl of a gadolinium-based contrast agent in 1×PBS (Gibco #10010023). After perfusion, the skull was removed with the brain intact within the skull to maintain anatomical integrity, and cranial bones were cleaned. The specimen was immersed in 30 ml of post-fixative solution containing 2% PFA and 120 µl gadolinium contrast agent in PBS and incubated overnight at room temperature. Samples were then transferred to storage solution (120 µl gadolinium contrast agent in 30 ml PBS) and kept at 4°C in sealed vials until imaging. For DiOlistic labelling: Following perfusion, brains were dissected and immersed in 4% PFA in PBS at 4°C for 30 min, washed 2-3 times in cold PBS for 1-2 h, and cryoprotected by sequential incubation in 15% sucrose (20 min, 4°C) and 30% sucrose (20 min, 4°C; Sigma #S9378) in PBS. After cryoprotection, tissue was stored in PBS at 4°C until use.

### Immunohistochemistry

Organs were either cryoprotected in 30% sucrose, embedded in Tissue-Tek^®^ O.C.T. (#4483; Sakura Finetek) and frozen on dry ice, or embedded in 4% low-melting-point agarose (Sigma-Aldrich, #A9414) and mounted on the holder for sectioning. The tissue was then cut coronally in 14 µm-thin sections for olfactory bulb analysis and 40 µm-thick floating sections for frontal cortex analysis. Immunostaining was performed by incubating the sections with the antibody (see Appendix Table S2 for dilutions). Sections of the olfactory bulb were imaged on the confocal Spinning disk (GATACA/Nikon) X1 Yokogawa head microscope on Prime95B (Photometrics) camera using MetaMorph (Molecular Devices^®^) software. Sections of the frontal cortex were imaged on the confocal laser scanning Zeiss LSM 980 Airyscan 2 microscope using Zen blue software. The LSM plus processing was applying during acquisition using a dry 20× NA:0.8 objective (total thickness: 20 µm; 11 slices; 2 µm interval). Images were analysed and adjusted using Adobe Photoshop or Fiji (NIH)^96^.

### MRI Imaging

After fixation, dissected skulls devoid of skin and muscle were stored in a 1:250 mixture of 0.5 mol gadoteric acid (Dotarem^®^, Guerbet) in PBS for at least 72 h. For imaging, the skulls were immersed in the immiscible fluorocarbon oil (Fluorinert™, Merck F9755) and immobilised in 15-ml conical-bottom tubes. Acquisitions were performed on a 17.2 T (1H Larmor frequency = 730.2 MHz) Bruker Biospec preclinical scanner equipped with a 25 mm inner diameter quadrature birdcage volume coil (Rapid Biomedical). T2-weighted 3D images were acquired using a Rapid Acquisition with Enhanced Relaxation (RARE) spin-echo sequence using the following parameters: [resolution 40 µm^3^, repetition time (TR): 1500 ms, effective echo time (TEeff): 24 ms, RARE factor: 4, number of averages (NA): 2, field-of-view (FOV): 18 mm × 12 mm × 8 mm, matrix size: 450×300×200, acquisition time: 12 h 30 min] (2-month-old and 1-year-old) or [resolution 50 µm^3^, repetition time (TR): 1500 ms effective echo time (TEeff): 30 ms, RARE factor: 6, number of averages (NA): 2, field-of-view (FOV): 18 mm × 14 mm × 14 mm, matrix size: 360×280×280, acquisition time: 10 h 44 min] (2-year-old).

### MRI image analysis

MRI datasets (Fig. 2b; S3) were processed using Fiji and Imaris (Bitplane). As automatic registration was impossible due to important morphological differences between wild-type and brains from several knockout mice, we decided to apply blinded semi-automatic analysis of scans. DICOM files were imported into Fiji, resliced into isotropic stacks (Image > Stacks > Reslice), and brightness/contrast were adjusted to optimise anatomical boundaries. The Scale&Segment V2 and Voxel Tools V2 macros were used to enable semi-automated segmentation and volumetric quantification. Regions of interest (ROIs) corresponding to brain structures such as the hippocampus, frontal cortex, olfactory bulb, and cerebellum were manually traced every fourth slice using the “O” tool, with intermediate slices interpolated automatically and gaps filled (Edit > Selection > Fill ROI holes). ROIs were combined into label masks, colour-coded using the Rainbow tool, and analysed with the Voxel tool to extract their volumes. For visualisation purposes, skull removal was performed by tracing ROIs excluding bone every 20 slices, interpolating masks, applying thresholds (min=2, max=255), binarizing, and smoothing edges with two iterations of dilation. Processed stacks were converted to 16-bit, pixel value divided by 255, reconverted to 8-bit, and inspected in the 3D Viewer to remove residual non-brain tissue, yielding skull-free brain images. Labelled structures were combined with skull-free brain images using the Merge Channels tool to generate multi-channel datasets compatible with Imaris.

For final 3D reconstruction, datasets were imported into Imaris, where surface rendering was applied with pseudo-colours assigned to each structure. The Surface Rendering module automatically generated volumetric objects, enabling both quantitative analysis of regional volumes and qualitative visualisation of spatial relationships. Imaris further allowed interactive model manipulation, including rotation, sectioning, and highlighting of regions. All labelled image stacks, masks, and volumetric data files were archived in structured project directories, and exported data were used for representation

*Diolistic Labelling and dendritic spine density analysis*

Dendritic spines were visualised using DiOlistic labelling (Fig. 5a,c; S6a,b; S7a,b) as described before^72, 73^. Tungsten and gold particles were coated with the lipophilic tracer DiI. Briefly, 300 mg tungsten powder was suspended in 300 µl dichloromethane, and 13.5 mg DiI was dissolved in 450 µl dichloromethane (3 mg/100 µl). Equal volumes (100 µl each) of the tungsten suspension and DiI solution were mixed on a glass slide, dried, finely ground, resuspended in 3 ml distilled water, and sonicated for 30 min. Gold–DiI particles were prepared similarly using 30 mg of 1 µm gold particles (Bio-Rad, #1652263). Gene Gun cartridges were prepared by coating Tefzel tubing with 10 mg/ml polyvinylpyrrolidone (PVP; Sigma, #437190), loading the DiI-coated tungsten suspension, drying under nitrogen, cutting the tubing into 13 mm segments, and storing the cartridges at 4°C until use.

Coronal brain sections (250 µm) were prepared using a vibratome, washed in PBS, and sequentially incubated in 15% and 30% sucrose for 5 min each. DiI-coated tungsten particles were delivered into the slices using a helium-powered Gene Gun system (Bio-Rad) at 120 psi. Following labelling, slices were washed with PBS to remove residual particles and incubated in PBS for 30 min in the dark to allow dye diffusion. Sections were mounted in 0.5% n-propyl gallate in 90% glycerol/PBS and imaged the following day using a Leica Stellaris confocal microscope, with either the HC PL APO CS2 63x/1.40 Oil or the HC PL APO CS2 40x/1.10 Water objectives, with a white light laser (range 440-790 nm) set to 551 nm. Dendritic spines were detected manually, and spine density was quantified using the Neurolucida (MFB Neuroscience) software in at least three wild-type and *Ttll1^-/-^*brains for every age and brain area analysed.

### Olfaction test

The olfaction test (Fig. S8a) was conducted on young adult and old mice following the protocol described before^97^. Odour stimuli included non-social odours: almond (Selectarôme, #AM-0824), and banana extracts (Selectarôme, #BA-0125), and social odours were obtained from bedding of unfamiliar cages. Non-social odour solutions were freshly prepared on the day of testing, and social odours were collected from cages that had not been changed for at least 3 days. Testing materials included clean standard mouse cages with fresh bedding (46×23.5×20 cm), clean cage lids, cotton-tipped wooden applicators (6-inch), weigh boats (4.5×4.5 cm), and digital timers for recording trial and sniffing times.

Mice were first acclimated for 30 min in a clean testing cage, during which a clean dry cotton-tipped wooden applicator was placed into the cage lid to minimize novelty-driven exploration during the test. Odour presentation followed a fixed sequence: three consecutive trials of water (control), followed by almond, banana, and two social odours. Each trial lasted 2 min, with a 1 min inter-trial interval during which applicators were replaced. Applicators were mounted on the cage lid with the cotton end positioned ∼4 cm into the cage. During trials, mice were recorded from a distance of ∼2 m, and cumulative time spent sniffing was measured with a stopwatch. Sniffing was defined as orienting toward the swab with the nose within ∼2 cm of the applicator. After each trial, the applicator was replaced, and animals from the same home cage were tested sequentially, though none were returned to the home cage until testing of all cage-mates was complete.

### IntelliCage^®^ behavioural tests

All behavioural experiments were performed using the IntelliCage^®^ system (TSE Systems^76^; Fig. S8b). Each unit consisted of a large acrylic cage (55 cm wide × 38 cm deep × 21 cm high) equipped with four operant corners, each containing RFID antennas, water bottle access modules, LED light cues, and air-puff units. The system allows testing mice in their social group with minimal interference from the experimenter. Individual mice were identified with subcutaneously implanted RFID transponders (Trovan #ID100A), enabling continuous monitoring of visits, nose pokes, and drinking behaviour. Mice were maintained under a 12 h light/dark cycle with water and food ad libitum. The evaluation of cognitive functions in the IntelliCage^®^ is done by the automated recording of mouse behaviour in the active corners of the cage, where mice need to perform nose-pokes to get access to drinking water (Fig. S8b).

The programmes used for behaviour tests include changes in exercises, and record the numbers of corner visits, nose pokes and licks for each animal throughout the duration of the test (Fig. 6a). During the adaptation phase (5–7 days), mice were habituated to the IntelliCage^®^ environment, with all corners providing free water access for 2 days, followed by 3-5 days of nose poke-based access (nose poke adaptation). This phase minimised novelty-driven exploration and established baseline corner preferences and drinking patterns. The total number of visits was measured daily and averaged to assess the spontaneous activity, while the number of visits without nose pokes was averaged per day as an index of exploratory behaviour^98^.

During the progressive ratio test, which assesses goal-oriented motivation (24 h), the number of nose pokes required to open the door progressively increased after each series of eight rewarded visits (2, 3, 4, 5, 6, 7, 8, 10, 12, 16, 20, and 24 nose pokes), with the time windows adjusted accordingly. The test ended after 24 h or when animals reached the 24-nosepoke module. After a 1-day pause, long-term memory was assessed using a fixed ratio test (24 h), in which mice were re-exposed to the same nose poke requirement reached at the end of the progressive ratio phase (i.e., the highest requirement achieved by each animal). Retention was evaluated based on the number of visits required to obtain access to water^99, 100^.

Spatial learning was assessed using the place learning test over a 2-day acquisition phase. During this phase, only one corner provided access to water: nose pokes at the correct corner opened the door, whereas nose pokes at the other corners were unrewarded. The number of visits to each corner was recorded and divided into blocks of 50 visits. The percentage of visits to the correct corner was used as a measure of spatial learning performance^99^.

Motor impulsivity and inhibitory control were assessed using a nose poke inhibition task (24 h), in which access to water was delayed following the initial nose poke. Delay intervals progressively increased across sets of 10 visits (0.5, 1, 1.5, 3, 4.5, 6, 7.5, and 9 s), as signalled by a blue LED on the doors. Premature pokes triggered a yellow LED and prevented water access until the animal exited the corner. The maximum delay reached by each animal was quantified as a measure of inhibitory control^98^.

The data from the IntelliCages were processed using Analyzer software (IntelliCage Plus, version 3.3.5.0; NewBehavior, TSE) and subsequently analysed using GraphPad Prism version 8.3.0 (GraphPad Software, USA). Outliers were identified using Grubbs’ test, performed with the GraphPad Outlier Calculator, with statistical significance set at α=0.05. Statistical analyses were selected according to the data distribution and experimental design.

A one-way ANOVA was used to assess differences between genotypes, independently for each age group. For variables measured repeatedly within the same animals, a two-way ANOVA was performed, with genotype as the between-subject factor and the number of visits as the within-subject factor. When a significant main effect was detected, post-hoc pairwise comparisons were performed using Tukey’s test. Statistical significance was set at p<0.05. Prior power analyses were conducted using G*Power software to minimize the number of animals used while ensuring adequate statistical power. No animals were excluded from the experimental groups; however, the final sample size per group was slightly reduced in specific analyses due to predefined exclusion criteria and occasional missing or incomplete data.

### Image analysis for immunofluorescence staining

Fluorescence images (Fig. 4a,c,e; S5; S6) were acquired using either a confocal spinning disk confocal microscope or a confocal laser scanning Zeiss LSM 980 Airyscan 2 system under identical acquisition settings across all samples, and analysed using Fiji (NIH) supplemented with custom macros and the Cellpose segmentation algorithm^101^. For quantification of NeuN- and DAPI-positive cells in the olfactory bulb, images were processed by converting to 8-bit greyscale, thresholding, and manual annotation using the Cell Counter plugin. NeuN- and DAPI-positive cells were scored as neurons, and data were expressed as the proportion of NeuN-over DAPI-positive cells, with counts obtained from 10 glomeruli per animal. For parvalbumin (PV) interneuron density in the frontal cortex, images were acquired as total thickness of 20 µm with 11 slices at 2 µm intervals using a 20× NA:0.8 objective, and PV-positive cells were manually counted in Fiji within anatomically matched ROIs; values were expressed in cells/mm². Frontal cortex area measurements were obtained from coronal sections using the line and polygon tools in Fiji to delineate anatomical boundaries relative to the olfactory bulb, with ROI area calculated automatically and averaged across five matched sections per animal. For NeuN quantification in the frontal cortex, DAPI signals were pre-processed with a gaussian filter (radius 1) and background were subtracted (rolling ball = 100) and nuclei were detected with Cellpose (diameter = 120 pixels). Segmented nuclei were exported as ROIs with the MIC-MAQ plugin, and both morphological and fluorescence intensity parameters were extracted for NeuN and DAPI channels, allowing identification of NeuN-positive neurons (value 1700) relative to the total number of nuclei detected on the DAPI image. All images were analysed with identical acquisition and processing parameters across genotypes and experimental conditions to ensure reproducibility.

### Mouse lifespan experiments

38 *Tau^P301S^* and 20 *Tau^P301S^Ttll1^-/-^*animals were used in this study. Animals were observed throughout their lifespan and were sacrificed once they attained well-defined endpoints. The number of animals surviving at each time point was plotted using the Kaplan-Meier method.

### Recombinant Tau expression and purification

Fluorescently tagged Tau wild-type (HIS6-3C-mNeonGreen-Tau2N4R-3C-StrepTag) and Tau^P301S^ (HIS6-3C-mNeonGreen-Tau2N4R-mutP301S-3C-StrepTag) in pET-11 were expressed in *E. coli* BL21(DE3)-RIPL strain. The cells were grown at 30°C until OD_600_=0.5-0.6; protein expression was then induced by 0.1 mM IPTG, and the cells were grown overnight at 16°C. Bacteria (3-4 g) were lysed in 45 ml of lysis buffer (50 mM Tris pH 8.0, 300 mM NaCl, 2 mM β-mercapto ethanol, 0.5 µL Benzonase, and 1× Protease inhibitor cocktail), sonicated (5 min; On/Off: 2/4 s), and centrifuged (40,000×g; 30 min; 4°C). The soluble fraction was then subjected to NiNTA purification (washing buffer: 50 mM Tris pH 8.0, 300 mM NaCl, 2 mM β-mercapto ethanol, 20 mM Imidazole), protein was eluted from the column (elution buffer: 50 mM Tris pH 8.0, 300 mM NaCl, 2 mM bME, 300 mM Imidazole), subjected to Strep-Tactin XT purification (washing buffer: 50 mM Tris pH 8.0, 300 mM NaCl, 2 mM β-mercapto ethanol, 20 mM Imidazole), and eluted using BXT buffer (100 mM Tris pH 8.0, 150 mM NaCl, 1 mM EDTA, 50 mM Biotin). The eluted protein was incubated overnight with C3 protease in dialysis (dialysis buffer: 50 mM Tris, 300 mM NaCl, 1 mM DTT). The sample was loaded onto a NiNTA column, and flow-through was collected after 1 h incubation. The purified protein was concentrated using a 50K centrifugal filter tube (Amicon Ultra-4, Merck). Protein concentration was measured with a NanoDrop ND spectrophotometer (Thermo Scientific) at 280 nm absorbance. Proteins were flash-frozen in liquid nitrogen and stored at -80°C. All purification steps were performed at 4°C.

### Microtubule Assembly

GMPCPP microtubules were polymerised from ∼4 mg/ml mouse brain tubulin (wild-type or *Ttll1^-/-^*) for 2 h at 37°C in BRB80 (80 mM PIPES pH 6.9, 1 mM EGTA, 1mM MgCl_2_) supplemented with 1 mM MgCl_2_ and 1 mM GMPCPP (#NU-405, Jena Bioscience). The polymerised microtubules were centrifuged for 30 min at 18,000×g in a Microfuge 18 Centrifuge (Beckman Coulter). After centrifugation, the pellet was resuspended and kept in BRB80 at room temperature.

### TIRF experiments

For TIRF experiments, chambers were assembled by melting thin strips of parafilm in between two glass coverslips silanised with Hexamethyldisilazane (HMDS, #379212). The chambers were incubated with 20 µg/mL anti-β-tubulin antibodies (in PBS, #T7816, Sigma) for 10 min, followed by 1% Pluronic (F127 in BRB80, #P2443, Sigma) for at least 30 min. Total internal reflection fluorescence (TIRF) microscopy experiments were performed on an inverted microscope (Nikon TI2 E) with an H-TIRF module equipped with a Nikon Apo 60× NA 1.49 oil immersion objective and a PRIME BSI (Teledyne Photometrics) camera.

Microtubules were visualised using interference reflection microscopy (IRM, CoolLED) and fluorescent proteins with a 488-nm laser. The microscopes were controlled with Nikon NIS-Elements software (v.5.42). All experiments were performed at room temperature.

Wild-type and *Ttll1^-/-^* microtubules were consecutively immobilised on the coverslip surface, and their position was imaged. Unbound microtubules were washed away with BRB80 and chambers were pre-incubated with TIRF assay buffer (50 mM HEPES pH 7.4, 1 mM EGTA, 2 mM MgCl_2_, 75 mM KCl, 10 mM dithiothreitol, 0.02 mg/ml casein, 1 mM Mg-ATP, 20 mM D-glucose, 0.22 mg/ml glucose oxidase and 20 µg/ml catalase). mNeonGreen-Tau (wild type or Tau^P301S^) was diluted in TIRF assay buffer and added to the measurement chamber with at least four-fold the chamber volume. Fluorescent signal on the microtubules was measured after 5 min incubation.

Microscopy data were analysed using Fiji. mNeonGreen-Tau fluorescent intensities on microtubules were measured by drawing a line along the microtubule and measuring the mean intensity. For background subtraction, the same line was then moved to an area directly adjacent to the microtubule where no microtubule is present, and the mean of the background was then subtracted from the mean on the microtubule.

---

**Figure S1:**
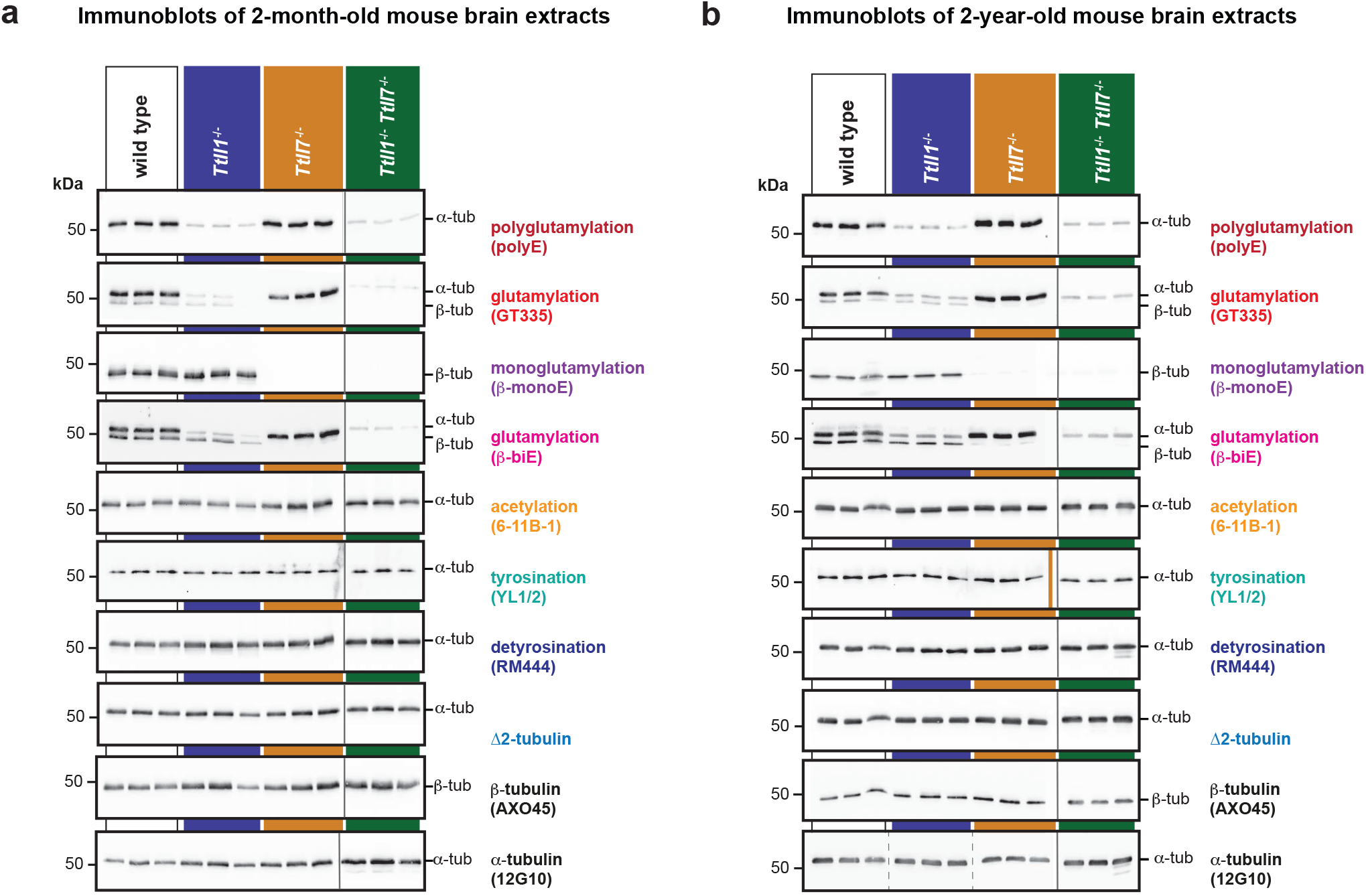
Characterisation of tubulin PTMs in brain extracts of mice at different ages. Immunoblot analysis of extracts of three mouse brains from 2-month-old wild-type, *Ttll1*^-/-^, *Ttll7*^-/-^, and *Ttll1*^-/-^*Ttll7*^-/-^ animals probed with polyE, GT335, β-monoE, and β-biE antibodies to detect different subtypes of (poly)glutamylation, antibodies for acetylation (6-11B-1), tyrosination (YL1/2), detyrosination (RM444), Δ2-tubulin. Total tubulin levels were determined with the pan-β-tubulin (AXO45) and pan-α-tubulin (12G10) antibodies. Molecular weight markers are indicated on the left (kDa), position of α- and β-tubulin bands are indicated on the right. Details of the antibodies are provided in Tables S1 and S2. Complementary to Fig. 1c.

**Figure S2:**
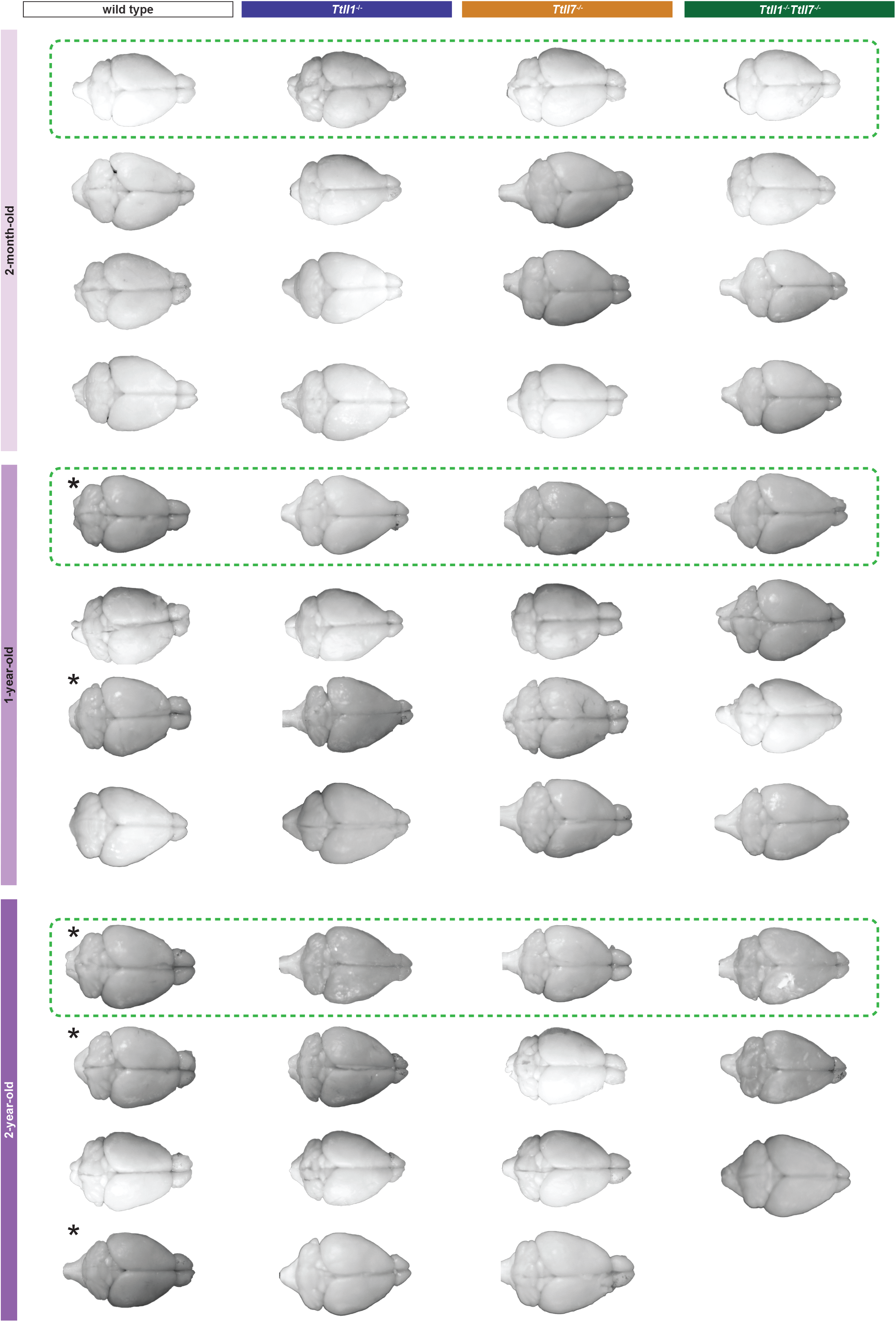
Macroscopic images of mouse brains across ages. Macroscopic images of brains from four (except *Ttll1*^-/-^*Ttll7*^-/-^ at 2 years) wild-type, *Ttll1*^-/-^, *Ttll7*^-/-^, and *Ttll1*^-/-^*Ttll7*^-/-^ mice at 2 months, 1 year, and 2 years. *denotes animals heterozygous for one of the genes. Images boxed in green dotted lines are also shown in Fig. 2a.

**Figure S3:**
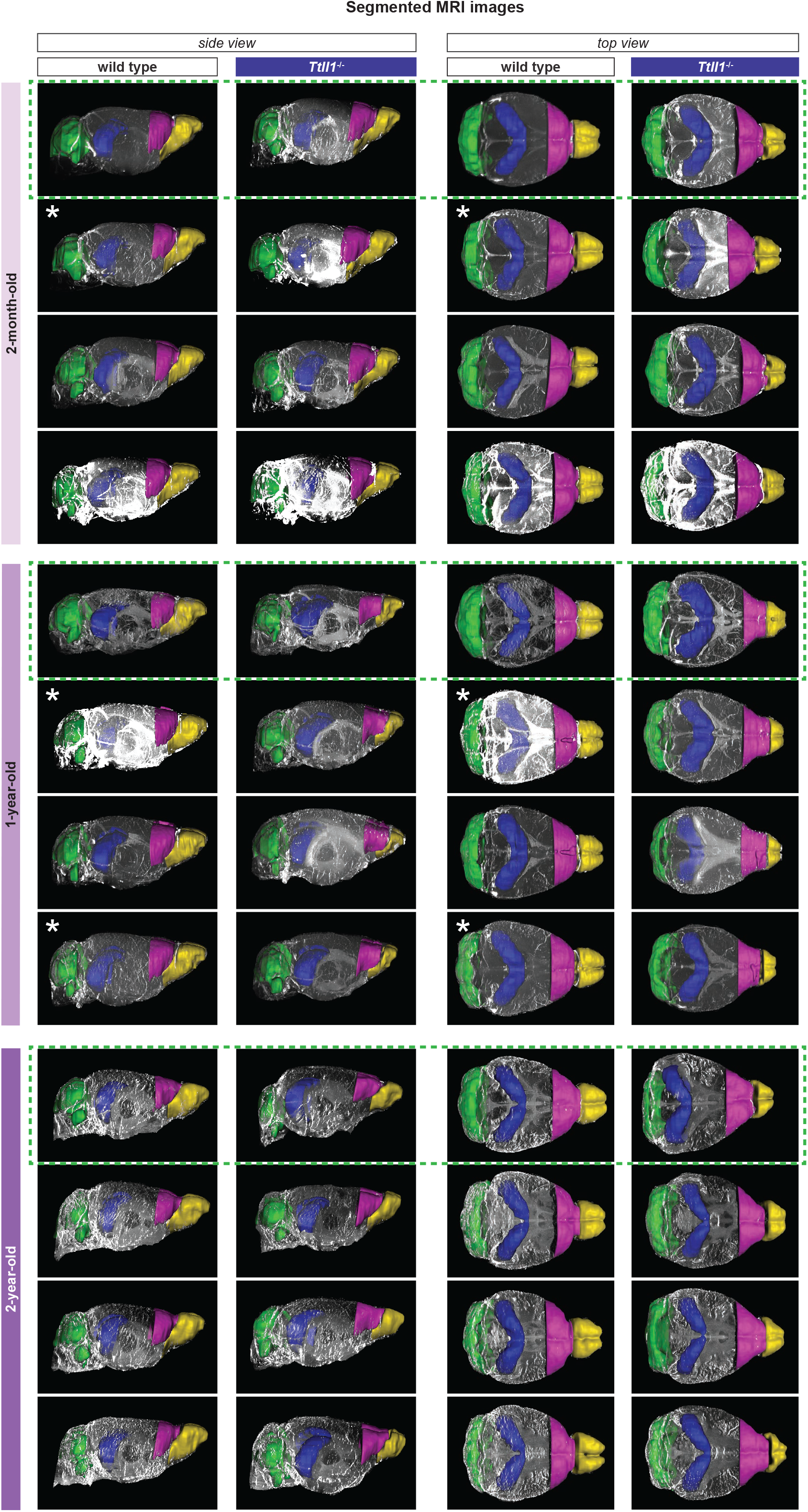
MRI-based brain segmentations of wild-type and *Ttll1*^-/-^ mice. Segmented three-dimensional reconstructions of MRI scans of brains from four wild-type and four *Ttll1*^-/-^ mice at 2 months, 1 year, and 2 years, displayed in side view (left panels) and top view (right panels) using Imaris. Different brain regions are color-coded: olfactory bulb (yellow), frontal cortex (pink), hippocampus (blue), and cerebellum (green). *denotes animals heterozygous for one of the gene. Images boxed in green dotted lines are also shown in Fig. 2b.

**Figure S4:**
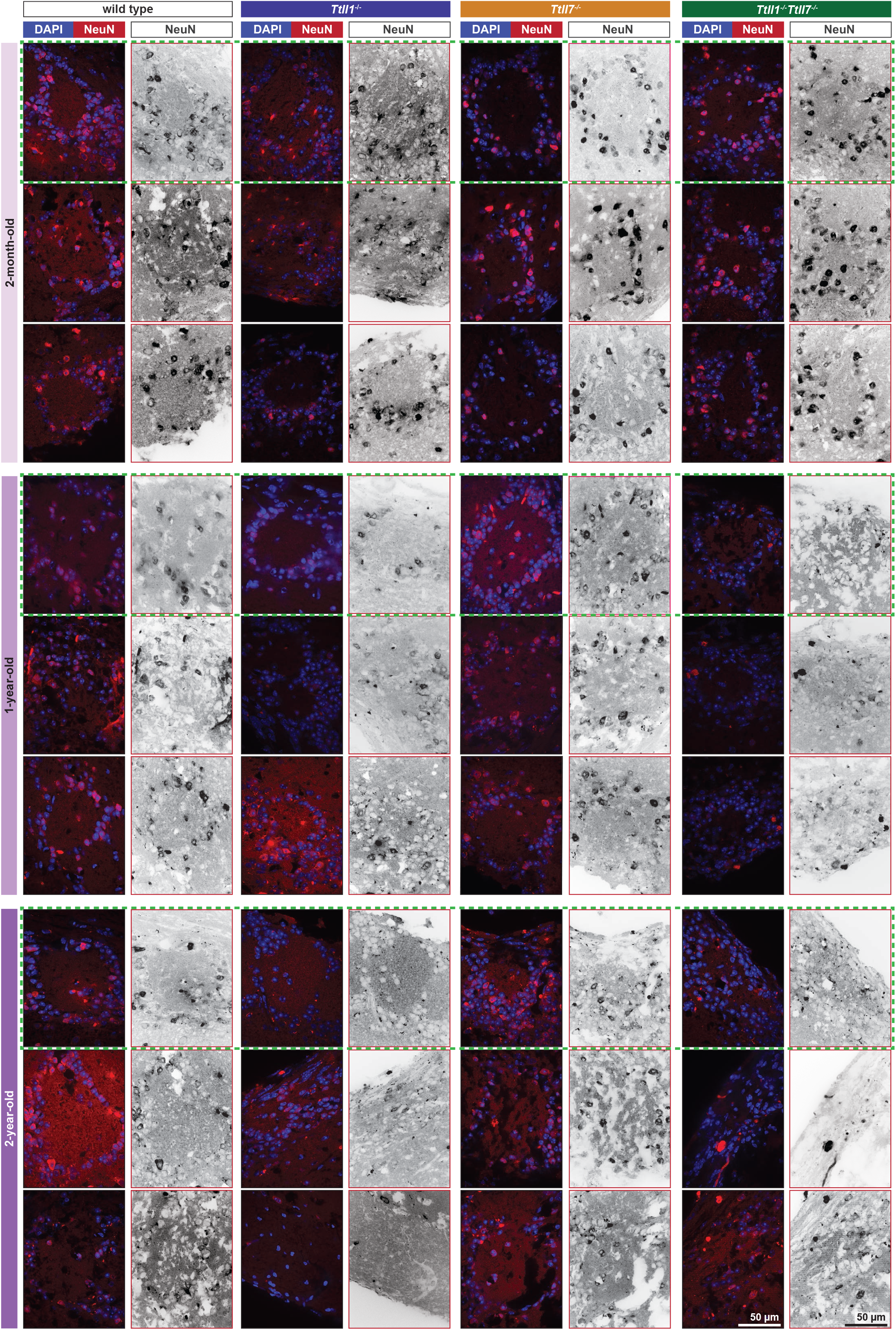
Immunohistological analyses of the olfactory bulb of mouse brains. Representative confocal images of NeuN immunostaining in the glomeruli of the olfactory bulb of three wild-type, *Ttll1*^-/-^, *Ttll7*^-/-^, and *Ttll1*^-/-^*Ttll7*^-/-^ mice at 2 months, 1 year, and 2 years. NeuN (red) labels nuclei of mature neurons, and DAPI (blue) marks all nuclei. Greyscale panels show NeuN single-channel images. Scale bar: 50 μm. * denotes animals heterozygous for one of the genes. Images boxed in green dotted lines are also shown in Fig. 3a.

**Figure S5:**
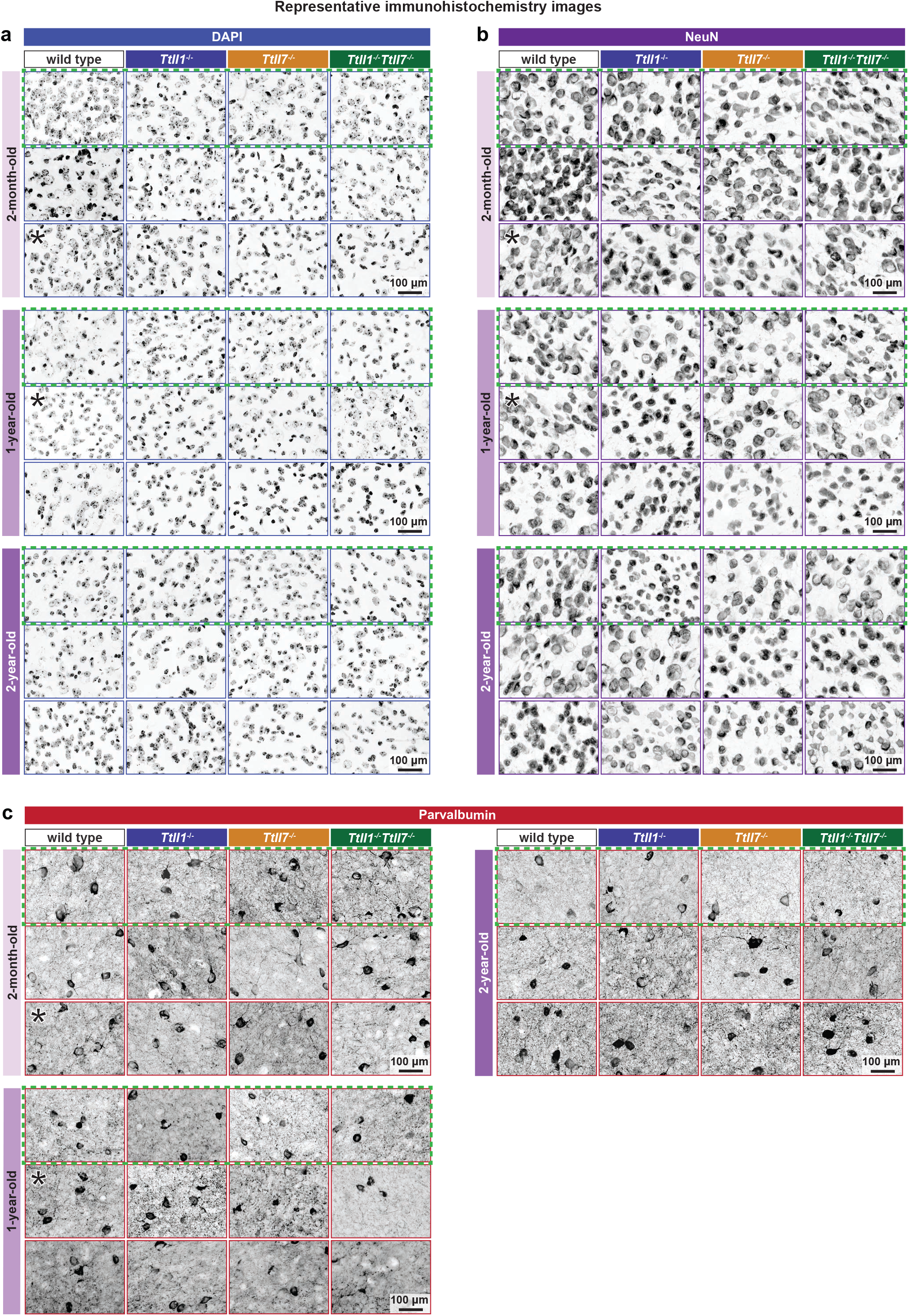
Immunohistological analyses of the frontal part of mouse brain cortex. Representative confocal images of **(a)** DAPI, **(b)** NeuN and **(c)** Parvalbumin (Pvn) staining used to determine overall cell densities (in cells/mm^2^) in the frontal part of the cortex of wild-type, *Ttll1*^-/-^, *Ttll7*^-/-^, and *Ttll1*^-/-^*Ttll7*^-/-^ brains at 2 months, 1 year, and 2 years (Fig. 4b,d,f). Scale bar: 100 μm. Each image corresponds to an individual animal. *denotes animals heterozygous for one of the genes. Images in green dotted boxes are also shown in Fig. 4a,c,e.

**Figure S6:**
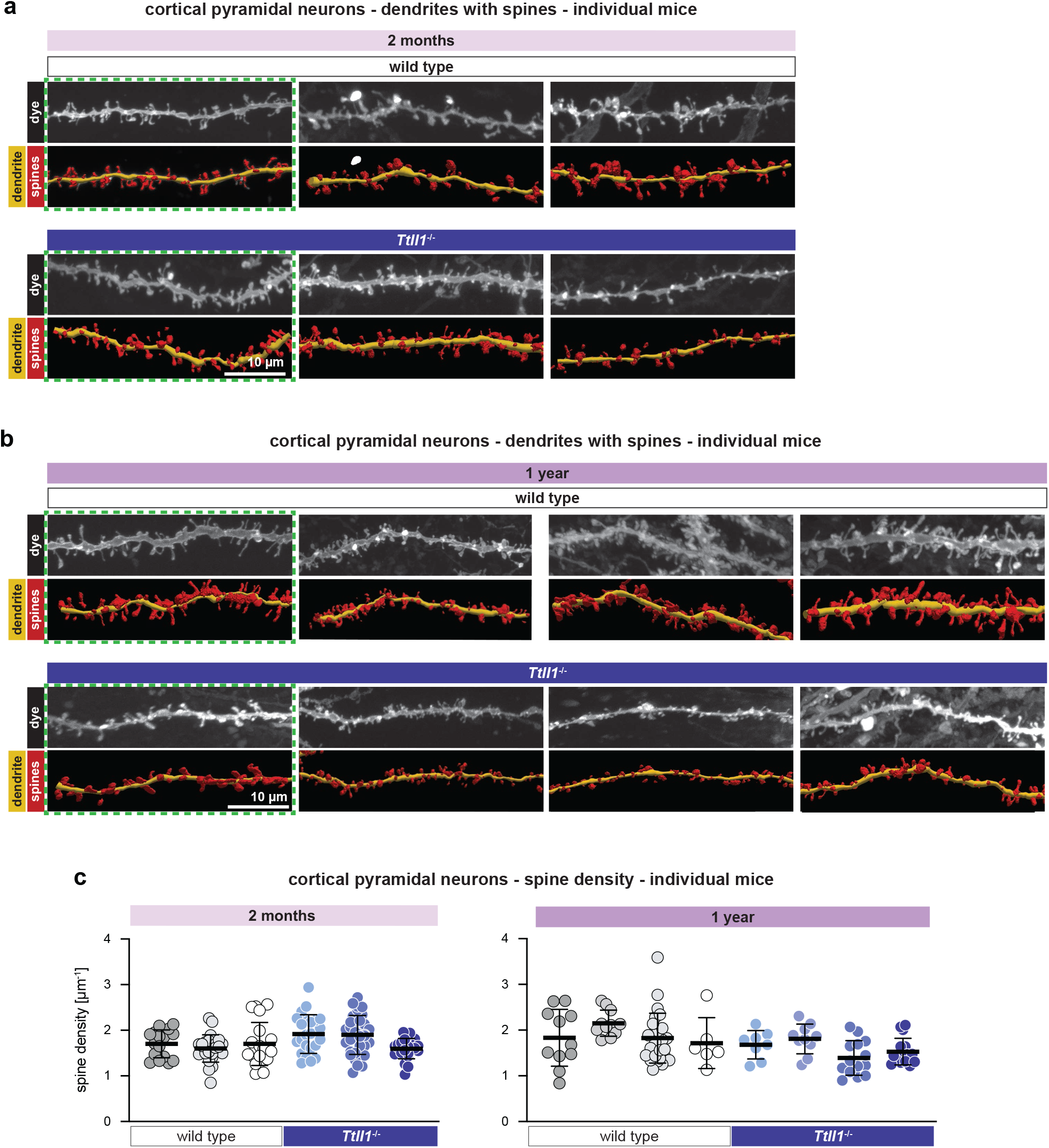
Dendritic spine density in cortical pyramidal neurons of wild-type and *Ttll1^-/-^*mice. **a, b)** Representative confocal images of dendritic segments from cortical pyramidal neurons of three 2-month-old (**a**) and tree 1-year-old (**b**) wild-type and *Ttll1*^-/-^ mice. Upper panels show greyscale dye-filled dendrites; lower panels show corresponding 3D reconstructions with dendrites in yellow and dendritic spines in red. Scale bar: 10 μm. Images in green dotted boxes are also shown in Fig. 5a. **c)** Quantification of dendritic spine densities (spines/μm) shown in (**a**) (2 months) and (**c**) (1 year). Each dot represents one analysed dendrite. Results obtained from different animals are shown in different colour shades (white/grey: wild type; blue: *Ttll1*^-/-^). Bars represent mean ± SD; fold differences are indicated. Combined analyses are shown in Fig. 5b.

**Figure S7:**
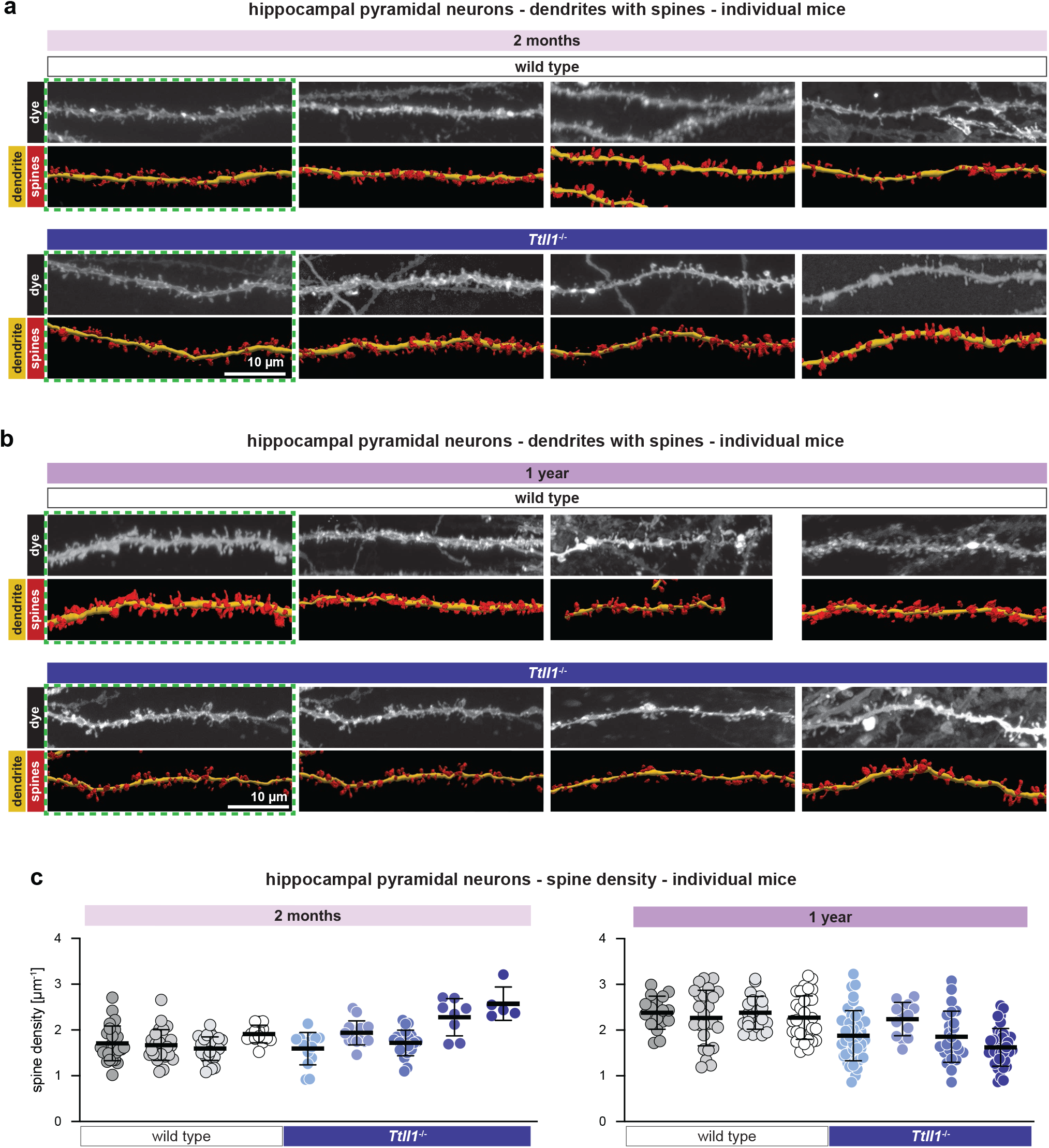
Dendritic spine density in hippocampal pyramidal neurons of wild-type and *Ttll1*^-/-^ mice. **a, b)** Representative confocal images of dendritic segments from hippocampal pyramidal neurons of four 2-month-old (**a**) and four 1-year-old (**b**) wild-type and *Ttll1*^-/-^ mice. Upper panels show greyscale dye-filled dendrites; lower panels show corresponding 3D reconstructions with dendrites in yellow and dendritic spines in red. Scale bar: 10 μm. Images in green dotted boxes are also shown in Fig. 5c. **c)** Quantification of dendritic spine densities (spines/μm) shown in (**a**) (2 months) and (**b**) (1 year). Each dot represents one analysed dendrite. Results obtained from different animals are shown in different colour shades (white/grey: wild type; blue: *Ttll1*^-/-^). Bars represent mean ± SD; fold differences are indicated. Combined analyses are shown in Fig. 5d.

**Figure S8:**
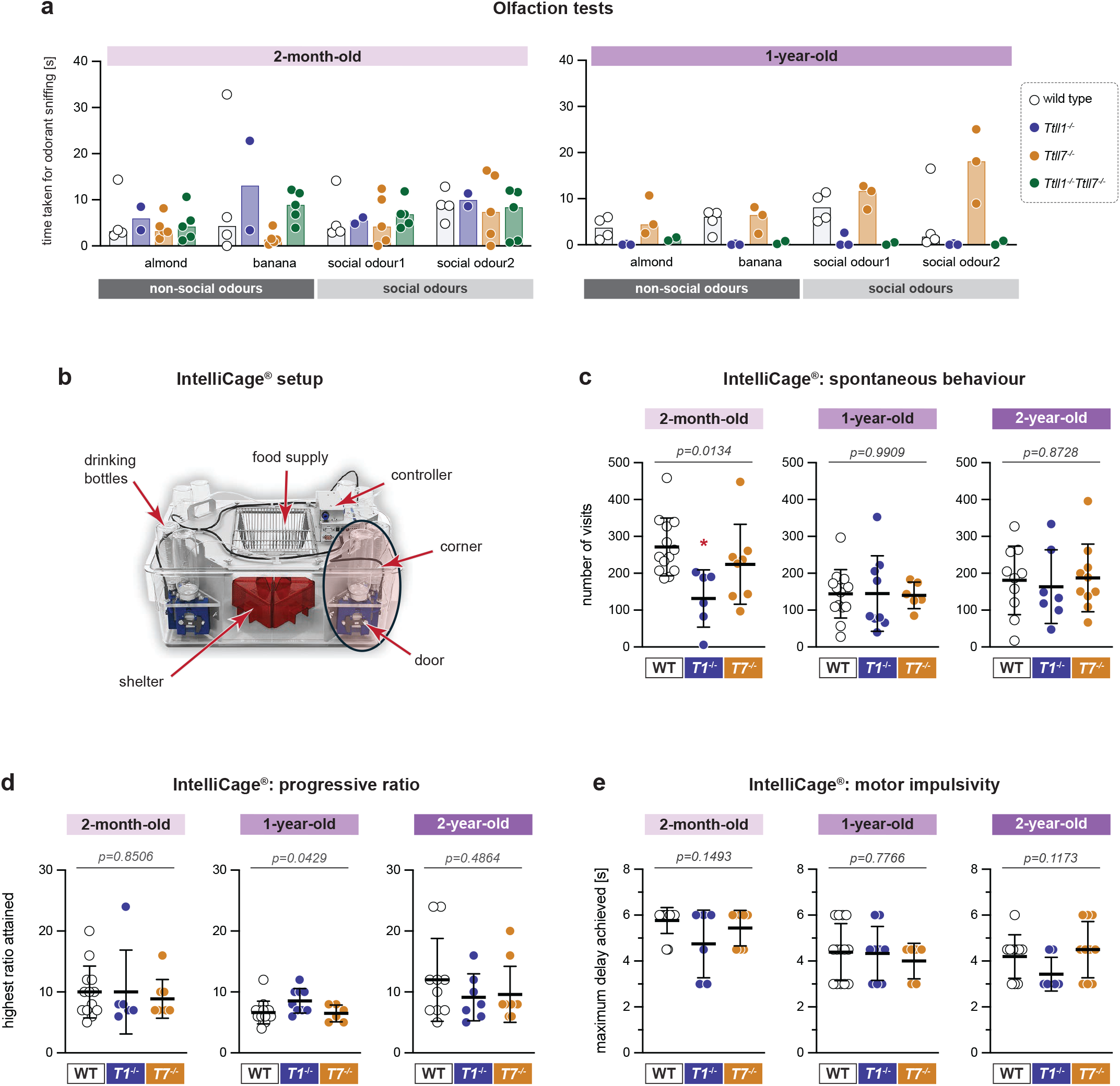
Behavioural effects of reduced polyglutamylation. **a)** Quantification of the olfaction test of 2-month-old and 1-year-old wild-type, *Ttll1*^-/-^, *Ttll7*^-/-^, and *Ttll1*^-/-^*Ttll7*^-/-^ mice. Time (in seconds) each animal took to sniff the odour is plotted. Each dot represents one animal; bars indicate median values. **b)** Illustration of the IntelliCage^®^ and its main components. **c, d, e)** Behavioural assessment carried out in the IntelliCage^®^ on 2-month-old, 1- and 2-year-old wild-type, *Ttll1*^-/-^ and *Ttll7*^-/-^ mice. Each dot represents one animal; bars represent mean ± SD. **c)** Assessment of spontaneous behaviour. Daily average of the visits to the corners in the first three days in the IntelliCage^®^. Higher values indicate higher spontaneous activity. Data were analysed using one-way ANOVA followed by Tukey’s post hoc test. *P* value indicates overall statistical significance of differences across genotypes. * p<0.05 *Ttll1^-/-^*vs. wild-type. **d)** Determination of learning capability (progressive ratio). The maximum number of nose pokes achieved (breakpoint) is plotted. Higher values indicate better learning capacities. Data were analysed using one-way ANOVA followed by Tukey’s post hoc test. *P* value indicates overall statistical significance of differences across genotypes. **e)** Quantification of motor impulsivity. The maximum delay time attained by each animal is plotted. Higher values indicate lower motor impulsivity. Data were analysed using one-way ANOVA. *P* value indicates overall statistical significance of differences across genotypes.

**Figure S9:**
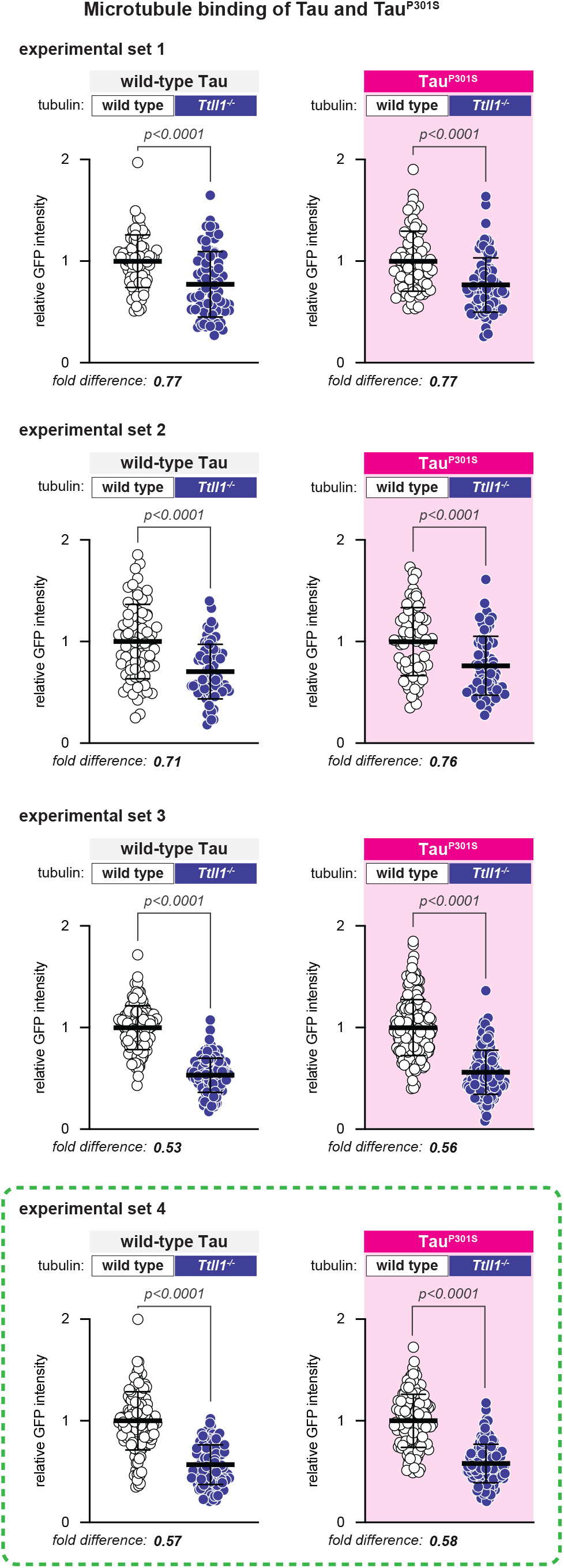
Quantification of wild-type Tau and Tau^P301S^ binding intensities. Quantification of four independent experiments showing wild-type Tau and Tau^P301S^ binding intensities to wild-type (open circles) and *Ttll1^-/-^* (blue). One experiment (boxed with green dotted line) is also shown in Fig. 7d. Fluorescence values were normalised to the mean value of wild-type microtubules in each experiment. Each data point represents a single microtubule. Bars represent mean ± SD. Fold differences are indicated. Statistical analyses were performed by two-tailed unpaired *t*-tests; *p* values are indicated.

**Table S1:**
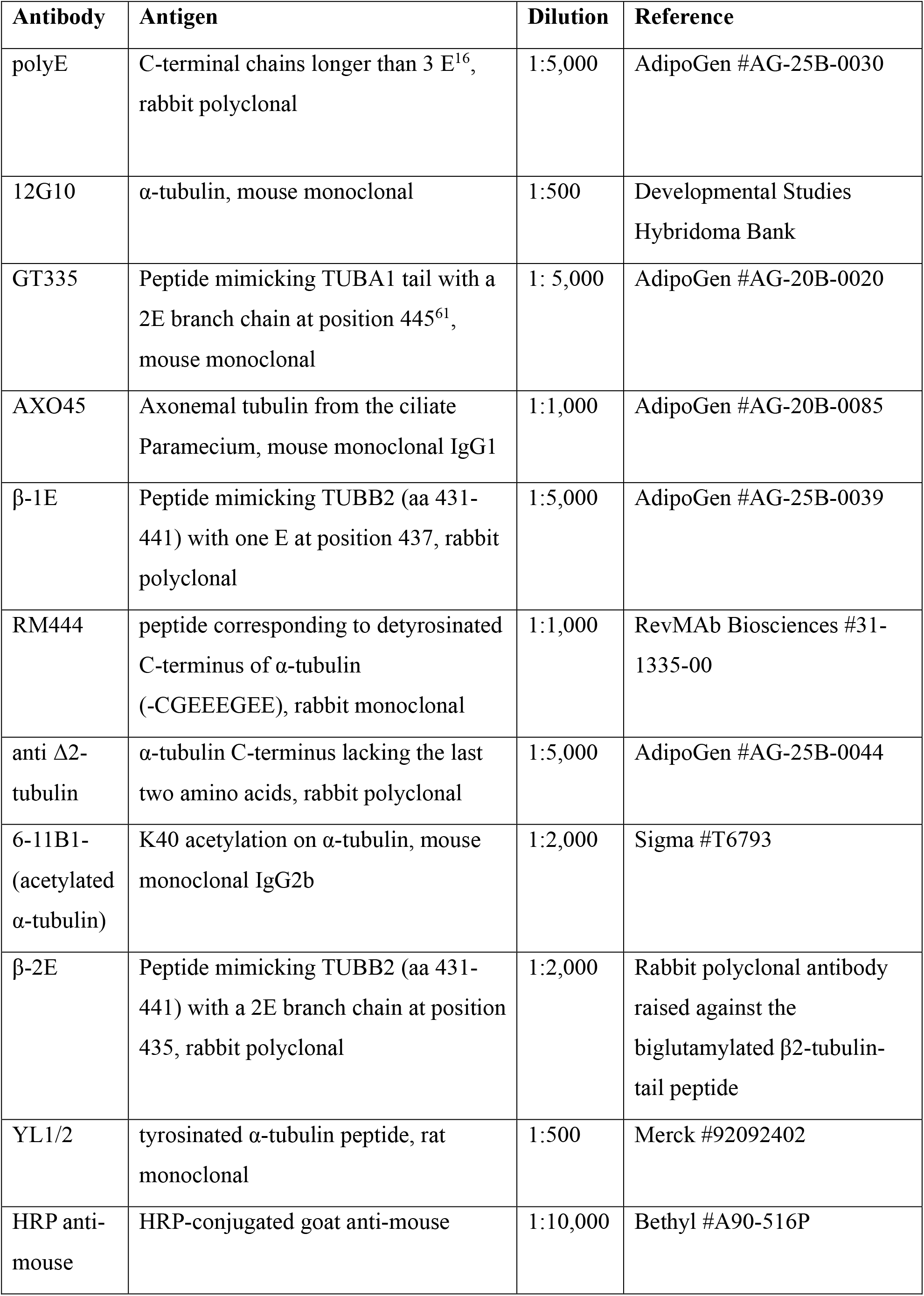
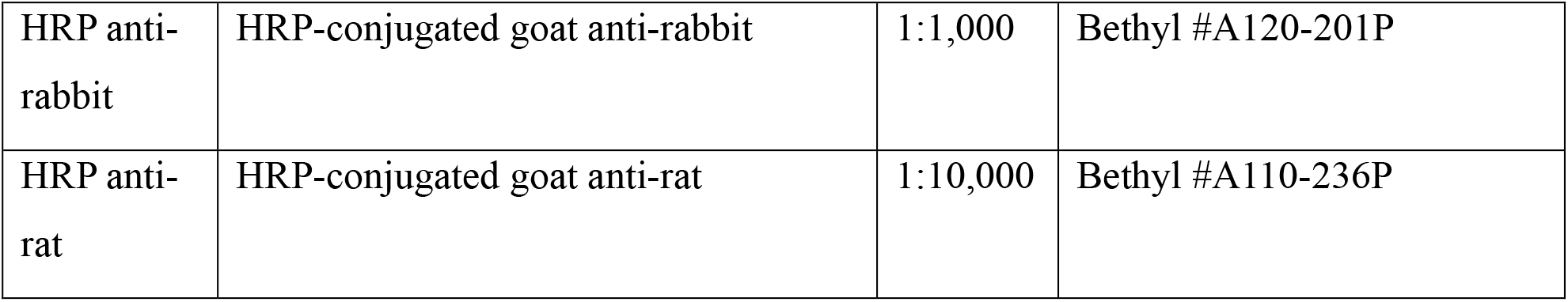
Primary and secondary antibodies used for immunoblotting.

| Antibody | Antigen | Dilution | Reference |
| --- | --- | --- | --- |
| polyE | C-terminal chains longer than 3 E<sup>16</sup>, rabbit polyclonal | 1:5,000 | AdipoGen #AG-25B-0030 |
| 12G10 | α-tubulin, mouse monoclonal | 1:500 | Developmental Studies Hybridoma Bank |
| GT335 | Peptide mimicking TUBA1 tail with a 2E branch chain at position 445<sup>61</sup>, mouse monoclonal | 1: 5,000 | AdipoGen #AG-20B-0020 |
| AXO45 | Axonemal tubulin from the ciliate Paramecium, mouse monoclonal IgG1 | 1:1,000 | AdipoGen #AG-20B-0085 |
| β-1E | Peptide mimicking TUBB2 (aa 431-441) with one E at position 437, rabbit polyclonal | 1:5,000 | AdipoGen #AG-25B-0039 |
| RM444 | peptide corresponding to detyrosinated C-terminus of α-tubulin (-CGEEEGEE), rabbit monoclonal | 1:1,000 | RevMAb Biosciences #31-1335-00 |
| anti Δ2-tubulin | α-tubulin C-terminus lacking the last two amino acids, rabbit polyclonal | 1:5,000 | AdipoGen #AG-25B-0044 |
| 6-11B1- (acetylated α-tubulin) | K40 acetylation on α-tubulin, mouse monoclonal IgG2b | 1:2,000 | Sigma #T6793 |
| β-2E | Peptide mimicking TUBB2 (aa 431-441) with a 2E branch chain at position 435, rabbit polyclonal | 1:2,000 | Rabbit polyclonal antibody raised against the biglutamylated β2-tubulin-tail peptide |
| YL1/2 | tyrosinated α-tubulin peptide, rat monoclonal | 1:500 | Merck #92092402 |
| HRP anti-mouse | HRP-conjugated goat anti-mouse | 1:10,000 | Bethyl #A90-516P |
| HRP anti-rabbit | HRP-conjugated goat anti-rabbit | 1:1,000 | Bethyl #A120-201P |
| HRP anti-rat | HRP-conjugated goat anti-rat | 1:10,000 | Bethyl #A110-236P |

**Table S2:** Primary and secondary antibodies used for immunofluorescence staining.

| **Antibody** | **Antigen** | **Dilution IF** | **Reference** |
| --- | --- | --- | --- |
| NeuN | NeuN | 1:200 | Abcam #177487 |
| Parvalbumin | Full length recombinant<br>rat parvalbumin | 1:500 | Synaptic Systems #195 004 |
| GFAP | Purified glial filament | 1:500 | Merck Millipore #MAB3402 |
| Alexa 488<br>anti-mouse | conjugated goat anti-mouse | 1:500-1:1,000 | Molecular probes #A11001 |
| Alexa 488<br>anti-rabbit | conjugated goat anti-rabbit | 1:500-1:1,000 | Molecular probes #A11008 |
| Alexa 568<br>anti-mouse | conjugated goat anti-mouse | 1:500-1:1,000 | Molecular probes #A11019 |
| Alexa 568<br>anti-rabbit | conjugated goat anti-rabbit | 1:500-1:1,000 | Molecular probes #A11036 |
| Alexa 647<br>anti-mouse | conjugated goat anti-mouse | 1:500-1:1,000 | Molecular probes #A21242 |
| Alexa 647<br>anti-rabbit | conjugated goat anti-rabbit | 1:500-1:1,000 | Molecular probes #A21245 |
| Alexa 568<br>anti-guinea<br>pig | conjugated goat anti-guinea pig | 1:500-1:1,000 | Invitrogen #A31931 |

## References

1. Atkins, M., Nicol, X. & Fassier, C. Microtubule remodelling as a driving force of axon guidance and pruning. Semin Cell Dev Biol 140, 35–53 (2023).

2. Miryala, C.S.J., Holland, E.D. & Dent, E.W. Contributions of microtubule dynamics and transport to presynaptic and postsynaptic functions. Mol Cell Neurosci 123, 103787 (2022).

3. Kelliher, M.T., Saunders, H.A. & Wildonger, J. Microtubule control of functional architecture in neurons. Curr Opin Neurobiol 57, 39–45 (2019).

4. Guedes-Dias, P. & Holzbaur, E.L.F. Axonal transport: Driving synaptic function. Science 366, science.aaw9997 (2019).

5. Sferra, A., Nicita, F. & Bertini, E. Microtubule Dysfunction: A Common Feature of Neurodegenerative Diseases. Int J Mol Sci 21, 7354 (2020).

6. Penazzi, L., Bakota, L. & Brandt, R. Microtubule Dynamics in Neuronal Development, Plasticity, and Neurodegeneration. Int Rev Cell Mol Biol 321, 89–169 (2016).

7. Brady, S.T. & Morfini, G.A. Regulation of motor proteins, axonal transport deficits and adult-onset neurodegenerative diseases. Neurobiol Dis 105, 273–282 (2017).

8. Sleigh, J.N., Rossor, A.M., Fellows, A.D., Tosolini, A.P. & Schiavo, G. Axonal transport and neurological disease. Nat Rev Neurol 15, 691–703 (2019).

9. Wethekam, L.C. & Moore, J.K. alphabeta-tubulin heterodimers: Origins and regulation of microtubule building blocks. Mol Biol Cell 37, re1 (2026).

10. Janke, C. & Magiera, M.M. The tubulin code and its role in controlling microtubule properties and functions. Nat Rev Mol Cell Biol 21, 307–326 (2020).

11. McKenna, E.D., Sarbanes, S.L., Cummings, S.W. & Roll-Mecak, A. The Tubulin Code, from Molecules to Health and Disease. Annu Rev Cell Dev Biol 39, 331–361 (2023).

12. Janke, C. et al. Tubulin polyglutamylase enzymes are members of the TTL domain protein family. Science 308, 1758–1762 (2005).

13. van Dijk, J. et al. A targeted multienzyme mechanism for selective microtubule polyglutamylation. Mol Cell 26, 437–448 (2007).

14. Rogowski, K. et al. Evolutionary divergence of enzymatic mechanisms for posttranslational polyglycylation. Cell 137, 1076–1087 (2009).

15. Wloga, D. et al. TTLL3 Is a tubulin glycine ligase that regulates the assembly of cilia. Dev Cell 16, 867–876 (2009).

16. Rogowski, K. et al. A family of protein-deglutamylating enzymes associated with neurodegeneration. Cell 143, 564–578 (2010).

17. Tort, O. et al. The cytosolic carboxypeptidases CCP2 and CCP3 catalyze posttranslational removal of acidic amino acids. Mol Biol Cell 25, 3017–3027 (2014).

18. Aillaud, C. et al. Vasohibins/SVBP are tubulin carboxypeptidases (TCPs) that regulate neuron differentiation. Science 358, 1448–1453 (2017).

19. Nieuwenhuis, J. et al. Vasohibins encode tubulin detyrosinating activity. Science 358, 1453–1456 (2017).

20. Landskron, L. et al. Posttranslational modification of microtubules by the MATCAP detyrosinase. Science 376, eabn6020 (2022).

21. Nicot, S. et al. A family of carboxypeptidases catalyzing alpha- and beta-tubulin tail processing and deglutamylation. Sci Adv 9, eadi7838 (2023).

22. Ersfeld, K. et al. Characterization of the tubulin-tyrosine ligase. J Cell Biol 120, 725–732 (1993).

23. Akella, J.S. et al. MEC-17 is an alpha-tubulin acetyltransferase. Nature 467, 218–222 (2010).

24. Shida, T., Cueva, J.G., Xu, Z., Goodman, M.B. & Nachury, M.V. The major alpha-tubulin K40 acetyltransferase alphaTAT1 promotes rapid ciliogenesis and efficient mechanosensation. Proc Natl Acad Sci U S A 107, 21517–21522 (2010).

25. Pagnamenta, A.T. et al. Defective tubulin detyrosination causes structural brain abnormalities with cognitive deficiency in humans and mice. Hum Mol Genet 28, 3391–3405 (2019).

26. Erck, C. et al. A vital role of tubulin-tyrosine-ligase for neuronal organization. Proc Natl Acad Sci U S A 102, 7853–7858 (2005).

27. Kalebic, N. et al. alphaTAT1 is the major alpha-tubulin acetyltransferase in mice. Nat Commun 4, 1962 (2013).

28. Ikegami, K. et al. Loss of alpha-tubulin polyglutamylation in ROSA22 mice is associated with abnormal targeting of KIF1A and modulated synaptic function. Proc Natl Acad Sci U S A 104, 3213–3218 (2007).

29. Magiera, M.M. et al. Excessive tubulin polyglutamylation causes neurodegeneration and perturbs neuronal transport. EMBO J 37, e100440 (2018).

30. Bosch Grau, M., et al. Tubulin glycylases and glutamylases have distinct functions in stabilization and motility of ependymal cilia. J Cell Biol 202, 441–451 (2013).

31. Gadadhar, S. et al. Tubulin glycylation controls axonemal dynein activity, flagellar beat, and male fertility. Science 371, eabd4914 (2021).

32. Eddé, B. et al. Posttranslational glutamylation of alpha-tubulin. Science 247, 83–85 (1990).

33. Rüdiger, M., Plessman, U., Kloppel, K.D., Wehland, J. & Weber, K. Class II tubulin, the major brain beta tubulin isotype is polyglutamylated on glutamic acid residue 435. FEBS Lett 308, 101–105 (1992).

34. Alexander, J.E. et al. Characterization of posttranslational modifications in neuron-specific class III beta-tubulin by mass spectrometry. Proc Natl Acad Sci U S A 88, 4685–4689 (1991).

35. Audebert, S. et al. Reversible polyglutamylation of alpha- and beta-tubulin and microtubule dynamics in mouse brain neurons. Mol Biol Cell 4, 615–626 (1993).

36. Audebert, S. et al. Developmental regulation of polyglutamylated alpha- and beta-tubulin in mouse brain neurons. J Cell Sci 107, 2313–2322 (1994).

37. Bodakuntla, S. et al. Tubulin polyglutamylation is a general traffic-control mechanism in hippocampal neurons. J Cell Sci 133, jcs241802 (2020).

38. Silva, C.G. et al. Cell-Intrinsic Control of Interneuron Migration Drives Cortical Morphogenesis. Cell 172, 1063–1078 (2018).

39. Chakraborty, S., Paulcan, S., Janke, C. & Magiera, M.M. Loss of Tubulin Tyrosination in Purkinje Neurons Does Not Cause Their Degeneration. bioRxiv, 2026.2007.2021.739800 (2026).

40. Wu, H.-Y. et al. TTLL1 and TTLL4 polyglutamylases are required for the neurodegenerative phenotypes in pcd mice. PLoS Genet 18, e1010144 (2022).

41. Bodakuntla, S. et al. Distinct roles of alpha- and beta-tubulin polyglutamylation in controlling axonal transport and in neurodegeneration. EMBO J 40, e108498 (2021).

42. Shashi, V. et al. Loss of tubulin deglutamylase CCP1 causes infantile-onset neurodegeneration. EMBO J 37, e100540 (2018).

43. Baltanas, F.C., Berciano, M.T., Santos, E. & Lafarga, M. The Childhood-Onset Neurodegeneration with Cerebellar Atrophy (CONDCA) Disease Caused by AGTPBP1 Gene Mutations: The Purkinje Cell Degeneration Mouse as an Animal Model for the Study of this Human Disease. Biomedicines 9, 1157 (2021).

44. Karakaya, M. et al. Biallelic variant in AGTPBP1 causes infantile lower motor neuron degeneration and cerebellar atrophy. Am J Med Genet A 179, 1580–1584 (2019).

45. Türay, S., Eröz, R. & Basak, A.N. A novel pathogenic variant in the 3’ end of the AGTPBP1 gene gives rise to neurodegeneration without cerebellar atrophy: an expansion of the disease phenotype? Neurogenetics 22, 127–132 (2021).

46. Sheffer, R. et al. Biallelic variants in AGTPBP1, involved in tubulin deglutamylation, are associated with cerebellar degeneration and motor neuropathy. Eur J Hum Genet 27, 1419–1426 (2019).

47. Gilmore-Hall, S. et al. CCP1 promotes mitochondrial fusion and motility to prevent Purkinje cell neuron loss in pcd mice. J Cell Biol 218, 206–219 (2019).

48. Krishnan, A. et al. Microtubule posttranslational modifications provide unique recognition patterns for associated proteins. EMBO J (2026).

49. Genova, M. et al. Tubulin polyglutamylation differentially regulates microtubule-interacting proteins. EMBO J 42, e112101 (2023).

50. Lacroix, B. et al. Tubulin polyglutamylation stimulates spastin-mediated microtubule severing. J Cell Biol 189, 945–954 (2010).

51. Valenstein, M.L. & Roll-Mecak, A. Graded Control of Microtubule Severing by Tubulin Glutamylation. Cell 164, 911–921 (2016).

52. Szczesna, E. et al. Combinatorial and antagonistic effects of tubulin glutamylation and glycylation on katanin microtubule severing. Dev Cell 57, 2497–2513 e2496 (2022).

53. Sotelo, C. Camillo Golgi and Santiago Ramon y Cajal: the anatomical organization of the cortex of the cerebellum. Can the neuron doctrine still support our actual knowledge on the cerebellar structural arrangement? Brain Res Rev 66, 16–34 (2011).

54. Hazan, J. et al. Spastin, a new AAA protein, is altered in the most frequent form of autosomal dominant spastic paraplegia. Nat Genet 23, 296–303 (1999).

55. Goedert, M., Eisenberg, D.S. & Crowther, R.A. Propagation of Tau Aggregates and Neurodegeneration. Annu Rev Neurosci 40, 189–210 (2017).

56. Chang, C.-W., Shao, E. & Mucke, L. Tau: Enabler of diverse brain disorders and target of rapidly evolving therapeutic strategies. Science 371, eabb8255 (2021).

57. Goedert, M. Alzheimer’s and Parkinson’s diseases: The prion concept in relation to assembled Abeta, tau, and alpha-synuclein. Science 349, 1255555 (2015).

58. Goedert, M., Crowther, R.A., Scheres, S.H.W. & Spillantini, M.G. Tau and neurodegeneration. Cytoskeleton (Hoboken) 81, 95–102 (2024).

59. Allen, B. et al. Abundant tau filaments and nonapoptotic neurodegeneration in transgenic mice expressing human P301S tau protein. J Neurosci 22, 9340–9351 (2002).

60. Magiera, M.M. & Janke, C. Investigating tubulin posttranslational modifications with specific antibodies, in Methods Cell Biol, Vol. 115, Edn. 2013/08/27. (eds. J.J. Correia & L. Wilson) 247–267 (Academic Press, Burlington; 2013).

61. Wolff, A. et al. Distribution of glutamylated alpha and beta-tubulin in mouse tissues using a specific monoclonal antibody, GT335. Eur J Cell Biol 59, 425-432 (1992).

62. Shang, Y., Li, B. & Gorovsky, M.A. Tetrahymena thermophila contains a conventional gamma-tubulin that is differentially required for the maintenance of different microtubule-organizing centers. J Cell Biol 158, 1195–1206 (2002).

63. Gant, M.S. et al. Unlocking the activities of polyglutamylation enzymes in brain tubulin with top-down mass spectrometry. ChemRxiv, 13 July 2026 (2026).

64. Nestor, S.M. et al. Ventricular enlargement as a possible measure of Alzheimer’s disease progression validated using the Alzheimer’s disease neuroimaging initiative database. Brain 131, 2443–2454 (2008).

65. Brinkman, S.D. & Largen, J.W., Jr. Changes in brain ventricular size with repeated CAT scans in suspected Alzheimer’s disease. Am J Psychiatry 141, 81–83 (1984).

66. Roy, D.S. et al. Memory retrieval by activating engram cells in mouse models of early Alzheimer’s disease. Nature 531, 508–512 (2016).

67. Whitman, M.C. & Greer, C.A. Adult neurogenesis and the olfactory system. Prog Neurobiol 89, 162–175 (2009).

68. Nagayama, S., Homma, R. & Imamura, F. Neuronal organization of olfactory bulb circuits. Front Neural Circuits 8, 98 (2014).

69. Kosaka, T. & Kosaka, K. Two types of tyrosine hydroxylase positive GABAergic juxtaglomerular neurons in the mouse main olfactory bulb are different in their time of origin. Neurosci Res 64, 436–441 (2009).

70. DeKosky, S.T. & Scheff, S.W. Synapse loss in frontal cortex biopsies in Alzheimer’s disease: correlation with cognitive severity. Ann Neurol 27, 457–464 (1990).

71. O’Brien, J.A. & Lummis, S.C.R. Diolistic labeling of neuronal cultures and intact tissue using a hand-held gene gun. Nat Protoc 1, 1517–1521 (2006).

72. Ziak, J. et al. CRMP2 mediates Sema3F-dependent axon pruning and dendritic spine remodeling. EMBO Rep 21, e48512 (2020).

73. Gavoci, A. et al. Polyglutamylation of microtubules drives neuronal remodeling. Nat Commun 16, 5384 (2025).

74. Dumont, M. Behavioral phenotyping of mouse models of neurodegeneration. Methods Mol Biol 793, 229–237 (2011).

75. Zou, J., Wang, W., Pan, Y.-W., Lu, S. & Xia, Z. Methods to measure olfactory behavior in mice. Curr Protoc Toxicol 63, 11 18 11–11 18 21 (2015).

76. Kiryk, A. et al. IntelliCage as a tool for measuring mouse behavior - 20 years perspective. Behav Brain Res 388, 112620 (2020).

77. Parra Bravo, C., Naguib, S.A. & Gan, L. Cellular and pathological functions of tau. Nat Rev Mol Cell Biol 25, 845-864 (2024).

78. Bugiani, O. et al. Frontotemporal dementia and corticobasal degeneration in a family with a P301S mutation in tau. J Neuropathol Exp Neurol 58, 667–677 (1999).

79. Souphron, J. et al. Purification of tubulin with controlled post-translational modifications by polymerization–depolymerization cycles. Nat Protoc 14, 1634–1660 (2019).

80. Berth, S.H. & Lloyd, T.E. Disruption of axonal transport in neurodegeneration. J Clin Invest 133, e168554 (2023).

81. Hosseini, S., van Ham, M., Erck, C., Korte, M. & Michaelsen-Preusse, K. The role of alpha-tubulin tyrosination in controlling the structure and function of hippocampal neurons. Front Mol Neurosci 15, 931859 (2022).

82. Ricci, C. Study on Genotypes and Phenotypes of Neurodegenerative Diseases. Genes (Basel) 15 (2024).

83. Kwok, J.B., Loy, C.T., Dobson-Stone, C. & Halliday, G.M. The complex relationship between genotype, pathology and phenotype in familial dementia. Neurobiol Dis 145, 105082 (2020).

84. Dejanovic, B., Sheng, M. & Hanson, J.E. Targeting synapse function and loss for treatment of neurodegenerative diseases. Nat Rev Drug Discov 23, 23–42 (2024).

85. Henstridge, C.M., Tzioras, M. & Paolicelli, R.C. Glial Contribution to Excitatory and Inhibitory Synapse Loss in Neurodegeneration. Front Cell Neurosci 13, 63 (2019).

86. Aiken, J. & Holzbaur, E.L.F. Cytoskeletal regulation guides neuronal trafficking to effectively supply the synapse. Curr Biol 31, R633–R650 (2021).

87. Hoffmann, N.A., Dorostkar, M.M., Blumenstock, S., Goedert, M. & Herms, J. Impaired plasticity of cortical dendritic spines in P301S tau transgenic mice. Acta Neuropathol Commun 1, 82 (2013).

88. Samaey, C., Schreurs, A., Stroobants, S. & Balschun, D. Early Cognitive and Behavioral Deficits in Mouse Models for Tauopathy and Alzheimer’s Disease. Front Aging Neurosci 11, 335 (2019).

89. Yoshiyama, Y. et al. Synapse loss and microglial activation precede tangles in a P301S tauopathy mouse model. Neuron 53, 337–351 (2007).

90. Adaikkan, C. et al. Alterations in a cross-hemispheric circuit associates with novelty discrimination deficits in mouse models of neurodegeneration. Neuron 110, 3091–3105 e3099 (2022).

91. Watamura, N. et al. In vivo hyperphosphorylation of tau is associated with synaptic loss and behavioral abnormalities in the absence of tau seeds. Nat Neurosci 28, 293–307 (2025).

92. Takeuchi, H. et al. P301S mutant human tau transgenic mice manifest early symptoms of human tauopathies with dementia and altered sensorimotor gating. PLoS One 6, e21050 (2011).

93. Hausrat, T.J. et al. Disruption of tubulin-alpha4a polyglutamylation prevents aggregation of hyper-phosphorylated tau and microglia activation in mice. Nat Commun 13, 4192 (2022).

94. Buscaglia, G., Northington, K.R., Moore, J.K. & Bates, E.A. Reduced TUBA1A Tubulin Causes Defects in Trafficking and Impaired Adult Motor Behavior. eNeuro 7, ENEURO.0045-0020.2020 (2020).

95. Lallemand, Y., Luria, V., Haffner-Krausz, R. & Lonai, P. Maternally expressed PGK-Cre transgene as a tool for early and uniform activation of the Cre site-specific recombinase. Transgenic Res 7, 105–112 (1998).

96. Schindelin, J. et al. Fiji: an open-source platform for biological-image analysis. Nat Methods 9, 676–682 (2012).

97. Yang, M. & Crawley, J.N. Simple behavioral assessment of mouse olfaction. Curr Protoc Neurosci Chapter 8, Unit 8 24 (2009).

98. Cisbani, G. et al. The Intellicage system provides a reproducible and standardized method to assess behavioral changes in cuprizone-induced demyelination mouse model. Behav Brain Res 400, 113039 (2021).

99. Poggini, S. et al. Minocycline treatment improves cognitive and functional plasticity in a preclinical mouse model of major depressive disorder. Behav Brain Res 441, 114295 (2023).

100. Poggini, S. et al. Combined Fluoxetine and Metformin Treatment Potentiates Antidepressant Efficacy Increasing IGF2 Expression in the Dorsal Hippocampus. Neural Plast 2019, 4651031 (2019).

101. Stringer, C., Wang, T., Michaelos, M. & Pachitariu, M. Cellpose: a generalist algorithm for cellular segmentation. Nat Methods 18, 100–106 (2021).

